# Protein Targets of Inositol Pyrophosphate (5-IP_7_) in the parasite *Trypanosoma cruzi*

**DOI:** 10.1101/2020.01.09.900860

**Authors:** Brian S. Mantilla, Nathaniel Brown, Dorothea Fiedler, Roberto Docampo

## Abstract

Diphosphoinositol-5-pentakisphosphate (5-PP-P_5_), also known as inositol heptakisphosphate (5-IP_7_), has been described as a high-energy phosphate metabolite that participates in the regulation of multiple cellular processes through protein binding or serine pyrophosphorylation, a post-translational modification involving a *β*-phosphoryl transfer. In this study, utilizing an immobilized 5-IP_7_ affinity reagent, we performed pull-down experiments coupled with mass spectrometry identification, and bioinformatic analysis, to reveal 5-IP_7_-regulated processes in the two proliferative stages of the unicellular parasite *Trypanosoma cruzi*. Our protein screen clearly defined two cohorts of putative targets either in the presence of magnesium ions or in metal-free conditions. The highest number of protein targets was detected in the absence of metal ions, with an overrepresentation of proteins involved in purine metabolism, protein phosphorylation, nucleosome assembly, cell redox homeostasis, parasite-surface components, and exocytosis, and only a 37% overlap between the two proliferative stages. In the presence of magnesium, there was a lower enrichment with the isolation of about 30 proteins with overrepresented candidates involved in tRNA amino acylation, phosphate homeostasis, lipid and glucose catabolism, and cell division, and 15% overlap between proliferative stages. To validate the pull-down results, we endogenously tagged four protein candidates and immunopurified them. Among the most interesting targets, we identified a choline/*o*-acetyltransferase domain-containing phosphoprotein that undergoes 5-IP_7_-mediated phosphorylation events at a polyserine tract (Ser^578-580^). We also identified a novel SPX domain-containing phosphoribosyl transferase [EC 2.7.6.1] herein termed as TcPRPS4. Our data revealed new functional roles of 5-IP_7_ in this divergent eukaryote, and provided potential new targets for chemotherapy.

## INTRODUCTION

Trypanosomatids are unicellular eukaryotes that can infect animals and humans causing deadly diseases. These parasites sense and respond to environmental stimuli to progress through their life cycle (1). *Trypanosoma cruzi* is a parasite transmitted by triatomine insects to vertebrates, including humans, in which they cause Chagas disease. Within its invertebrate host *T. cruzi* proliferates as epimastigote form while in the cytoplasm of the mammalian host it replicates as amastigote form. There are morphological, metabolic, and physiological differences between both proliferative stages of *T. cruzi* that clearly define them as distinct cells. In this sense, the understanding of signal transduction pathways and their interaction with metabolic networks is needed to define essential mechanisms for survival in its different hosts (2).

*Myo*-inositol phosphates (IPs) are a class of phosphorylated metabolites involved in the regulation of several cellular processes through their interaction with specific protein substrates. They are ubiquitously present in animal and plant cells as soluble products, and their synthesis relies on the activity of specific kinases that use ATP as phosphate donor (3). The canonical route for IP synthesis involves the action of phospholipase C on phosphatidylinositol 4,5-bisphosphate (PIP_2_) releasing inositol 1,4,5-trisphosphate (IP_3_) and diacylglycerol (4). IP_3_ has a well-studied receptor, the IP_3_ receptor, which participates in Ca^2+^ release from intracellular stores, endoplasmic reticulum (ER) in eukaryotes (5), and acidocalcisomes in trypanosomes (6). This signaling pathway was the first direct evidence of IP_s_-mediated processes through protein binding (5). IP_3_ can be further phosphorylated at different hydroxyl positions producing the fully phosphorylated form known as inositol hexakisphosphate (IP_6_) or phytic acid, which can be a precursor of the inositol pyrophosphates (7). Inositol pyrophosphates are characterized by the presence of single (PP-IP_4_ and PP-IP_5_) or double (PP_2_-IP_3_ and PP_2_-IP_4_) pyrophosphate moieties at different positions of the *myo*-inositol scaffold (8). In *Trypanosoma brucei*, the biosynthesis of 5-diphosphoinositol pentakisphosphate (5-PP-IP_5_ or 5-IP_7_) takes place in the cytosol by the action of IP6K using IP_6_ as substrate and ATP as phosphate donor (9). Once produced, 5-IP_7_ may interact with proteins at different cell compartments through two different mechanisms. One involves the transference of a *β*-phosphoryl group from 5-IP_7_ onto a phosphorylated serine surrounded by acidic residues. This posttranslational modification is Mg^2+^-dependent and was denominated as protein pyrophosphorylation (10). In the second mechanism, IP_7_ allosterically binds to a target protein through interaction with specific domains such as pleckstrin homology (PH) (3) or SPX domains (11). The SPX domains were initially found in three proteins from yeast, plants and human cells (Syg1, Pho81, XPR1) (12), but recent evidence in yeast has shown that the biosynthesis of polyphosphate is also regulated by 5-IP_7_ through its association to a SPX domain present in the vacuolar transporter chaperone (VTC) complex, which acts as a polyphosphate polymerase (11, 13). Molecular and physiological studies done with the phosphate/sodium symporters of *T. brucei* acidocalcisomes (TbPho91) and yeast vacuoles (Pho91p) demonstrated that 5-IP_7_ binds to their SPX domains and this is essential for phosphate release to the cytosol (14). Similarly, 5-IP_7_ binds to the SPX domain of the mammalian XPR1 (Xenotropic and Polytropic Retrovirus Receptor 1) phosphate transporter of the plasma membrane, promoting cellular phosphate export (15). Therefore, 5-IP_7_ has been proposed as a signaling molecule with critical importance in phosphate homeostasis and the cross talk between ATP, polyphosphate, and inositol pyrophosphates has relevant implications in eukaryotes (16).

In this work, we identified potential protein substrates for 5-IP_7_ in the two proliferative stages of *T. cruzi*. We uncovered a SPX-domain containing phosphoribosyl pyrophosphate synthetase that is unique and conserved across kinetoplastids. Also, we verified that 5-IP_7_-mediated phosphorylation is present in *T. cruzi*, and one of their targets contain a choline/carnitine *o*-acetyl transferase domain that could be involved in carbon metabolism in this pathogenic parasite.

## EXPERIMENTAL PROCEDURES

### Experimental Design and Statistical Rationale

Proteins pulled down from lysates of both *T. cruzi* proliferative stages were analysed by LC-MS/MS. Two independent biological experiments for each stage were done and the results of one of these experiments for each stage are shown. Only those proteins having a ≥ 4-fold change difference in their peptide-to-spectrum matches were considered for subsequent analysis. When indicated, values are expressed as means ± s.e.m. Significant differences between treatments were compared using the tests indicated in the figure legends. Differences were considered statistically significant at *P* < 0.05, and *n* refers to the number of independent biological experiments performed. All statistical analyses were conducted using GraphPad Prism 6 (GraphPad Software, San Diego, CA).

### Culture Methods

*Trypanosoma cruzi* epimastigotes (Y-strain, discrete-typed unit Tc-II) were maintained at 28°C in axenic cultures by successive passages in LIT medium (17) supplemented with 10% heat-inactivated fetal bovine serum (FBS) and 2 mg/ml of hemin. The epimastigotes transformed using the CRISPR/Cas9 system were maintained in the presence of 0.25 mg/ml geneticin (G418) and 0.15 mg/ml hygromycin (*HYG*). Tissue culture cell-derived trypomastigotes were collected from the supernatants of infected host cells, using a modification of the method of Schmatz and Murray (18), as previously described (19). For amastigote isolation, gamma-irradiated (exposed to 2,000 rads) Vero cells were seeded into 125 cm^2^ plastic flasks containing RPMI medium supplemented with 10% fresh FBS. After attachment, cell-derived trypomastigotes (collected at day 6 after infection) were added in a 10:1 ratio (parasites: host cell) and cells were incubated at 37°C under a 5% CO_2_ atmosphere for 4 h. Non-internalized parasites were removed by three washes with Hank’s balanced salt solution (HBSS) and RPMI-10% FBS was freshly added. Media was replaced daily and after ten days amastigotes could be collected from the culture supernatants. Detached host cells were removed by centrifugation (600 x *g* for 5 min) and the supernatant containing parasites was centrifuged (1,600 x *g* for 10 min). The parasite pellet was washed twice with HBSS, and centrifuged again. Two ml of fresh RPMI-10% FBS medium were added to the final pellet, centrifuged again, and then incubated for additional 2 h at 37°C under CO_2_. The supernatant was carefully removed with micropipette and amastigotes were resuspended in 500 µl of buffer A with glucose (BAG buffer: 116 mM NaCl, 5.4 mM KCl, 0.8 mM MgSO_4_, 50 mM Hepes-KOH, pH 7.2, and 5.5 mM D-glucose). Cell numbers were determined by counting on hemocytometer and pellets containing 2-4 x 10^9^ amastigotes were used for protein lysate preparation.

### Analysis of Total IP_6_ and 5-IP_7_ Content by LC-MS

Total inositol phosphates (IPs) were extracted from epimastigote and amastigote forms by adapting a method previously reported (20). Cell densities were determined prior to extraction and then used for data normalization. After harvesting the cells, pellets were stored at −80°C and thawed for extraction by adding 0.5 ml of 20% (w/v) freshly-prepared trichloroacetic acid (TCA) and mixing by vigorous vortexing. One volume of MS-grade ultrapure water (UP-water) was added to the samples, which were supplemented with 10 nmol of 3-deoxy-3-fluoro-D-*myo*-inositol 1,4,5-triphosphate, hexasodium salt (3F-IP_3_) (EMD Millipore), used as internal standard. Samples were vortexed and centrifuged at 12,000 x *g* for 20 min at 4°C. Supernatants were neutralized with 3% ammonium hydroxide and extracted with 2 ml of water-saturated diethyl ether (Sigma), vortexed and centrifuged at 2,000 x *g* for 10 min at 4°C. The organic phase was discarded and extraction was repeated one more time. The bottom aqueous phase was then loaded into a Strata™-X-AW 33µm polymeric weak anion column (Phenomenex®) preequilibrated with UP-water and methanol (MeOH, Merck) (1 ml each step). The column was washed with 25% MeOH and 100 mM ammonium acetate pH 7.2, and inositol phosphates were eluted by adding 1 ml of 100 mM ammonium carbonate (Sigma) and equilibrated with 0.5 mL of 5% acetic acid. Samples were dried with a speed vacuum, resuspended in 60 µl of UP-water and filtered by passing through nanoFilter™ 0.2 µm PVDF (Thomson) prior to LC-MS analysis. Ten µl of each sample were injected onto an Agilent 1100 HPLC system equipped with a BioBasic AX anion-exchange column (150 mm, ID 2.1 mm x 150 mm, 5 µm particle size) (Thermo) to separate IPs. Analytes were detected in a Quadrupole Time-of-Flight (ToF) mass spectrometer (Micromass). IPs were eluted by running a 45-min gradient of two mobile phases: buffer system A (25% MeOH: water) and system B (300 mM ammonium carbonate, pH: 9.0). The ESI source was set in negative-ion mode, with a spray voltage of 3,000 V, ion-transfer capillary T° at 300°C. Determination of total 5-IP_7_ content was extrapolated from standard curves ranging from 1-10 nmol of 5-IP_7_ (kindly provided by Dr. H. Jensen, University of Freiburg) and calculated using a variable number of epimastigotes (2 × 10^8^ −2 × 10^9^ parasites total). The cell volume of epimastigotes was considered as 34.8 ± 3.3 µl/10^9^ cells (21).

### Affinity Enrichment of 5-IP_7_ Substrates

For protein enrichment experiments samples from both stages were prepared in the presence of magnesium ions (Mg^2+^) or under metal-free conditions (EDTA). A 120-175 µL suspension of affinity reagents 5PCP-IP_5_ (a 5-IP_7_ non-hydrolyzable analog) or C (phosphate assay control) were prepared to result in approximately 120 nmol of each reagent on beads, as described previously (22). Briefly, beads were washed three times with binding buffer (50 mM Tris-HCl, pH 7.5, 150 mM NaCl, 0.05% Triton X-100), supplemented with 1 mM EDTA or 1 mM MgCl_2_. Then, total protein lysates from epimastigote and amastigote forms were suspended in lysis buffer [50 mM Tris-HCl, pH 7.5, 150 mM NaCl, 0.1% Triton X-100, 1X cOmplete™ EDTA-free protease inhibitor, PhosSTOP™ phosphatase inhibitor mixture (Roche) and 1 mM phenylmethanesulfonyl fluoride (PMSF)] supplemented with 1 mM EDTA or 1 mM MgCl_2_. Samples were kept on ice for 15 min and then lysed by sonication (3 cycles of 2 sec each at 15% amplitude/10 s on ice). Lysates were clarified by centrifugation (12,000 x *g* for 20 min at 4°C) and the supernatant was kept on ice. Protein concentration was determined by using the bicinchoninic acid (BCA) assay following the manufacturer indications (Thermo). Then, 1 mg of total protein from each lysate was incubated with the beads for 2 h, at 4°C, under gentle agitation. After this, beads were washed with 1 mL of binding buffer containing either MgCl_2_ or EDTA and centrifuged at 600 x *g* for 3 min. Elution of proteins was done by adding 100 µl of binding buffer supplemented with 20 mM IP_6_-sodium salt (Sigma) and then incubating overnight at 4°C under constant mixing. An additional elution step was done adding 50µL of elution buffer and incubating for 15 min.

### Protein Mass Spectrometry Data Analysis

The gel pieces were sliced into small pieces and then rinsed twice with 50% acetonitrile/20 mM ammonium bicarbonate (∼pH 7.5-8). Then, samples were dehydrated by adding 100% acetonitrile and dried out by speed vacuum. A various amount of Trypsin solution (0.01 µg/µl in 20 mM ammonium bicarbonate) was added until the gel pieces totally absorbed the Trypsin solution. The tubes were placed in an incubator at 37°C overnight. The tryptic peptides were extracted from gel pieces by incubating twice with 50% acetonitrile/0.1% formic acid. The extracts were dried down using a SpeedVac. The mass spectrometry analyses were performed on a Thermo-Fisher LTQ Orbitrap Elite Mass Spectrometer coupled with a Proxeon Easy NanoLC system (Waltham, MA) located at Proteomics and Mass Spectrometry Facility, University of Georgia. The enzymatic peptides were loaded onto a reversed-phase column (self-packed column/emitter with 200 Å 5 µM Bruker MagicAQ C18 resin), then directly eluted into the mass spectrometer. Briefly, the two-buffer gradient elution (0.1% formic acid as buffer A and 99.9% acetonitrile with 0.1% formic acid as buffer B) starts with 5% B, holds at 5% B for 2 min, then increases to 25% B for 60 min, to 40% B in 10 min, and to 95% B for 10 min. The data-dependent acquisition (DDA) method was used to acquire MS data. A survey MS scan was acquired first, and then the top 5 ions in the MS scan were selected for following CID and HCD MS/MS analysis. Both MS and MS/MS scans were acquired by Orbitrap at the resolutions of 120,000 and 30,000, respectively. Data were acquired using Xcalibur software (version 2.2, Thermo Fisher Scientific). Proteins identification and modification characterization were performed using Thermo Proteome Discoverer (version 1.4) with Mascot (Matrix Science) and TriTrypDB database. The spectra of possible modified peptides were inspected further to verify the accuracy of the assignments. The semi-quantitative analyses were achieved using a label-free quantification workflow within Proteome Discoverer, which a precursor ion areas detector node calculates the average of the top three highest peptide areas in the Extracted Ion Chromatograms (mass precision, 2 ppm). Positive hits were considered based on the peptide-spectrum match (PSM) obtained from 5PCP-IP_5_ enrichment and compared to the one from phosphate control beads. Only those proteins having a ≥ 4-fold change difference were considered for subsequent analysis. For bioinformatic analysis, protein sequences were retrieved from TriTrypDB database (tritrypdb.org) and then GO annotation workflow was run using the Blast2GO v5.2.1 software (23). Phosphoprotein analysis of identified candidates was made through a search of post-translational modifications with mass spectrometry evidence for phosphoproteomics with annotated entries in the TritrypDB. For comparative analysis of PRPSs among eukaryotes a phylogenetic reconstruction using protein sequences from distinct organisms was made. Protein search was made using BLAST tool with PRPSs from *T. cruzi* as query. Proteins were downloaded from UniprotKB (www.uniprot.org) or EupathDB (www.eupathDB.org) databases aligned and analyzed by the maximum likelihood method implemented in the PhyML program (v3.1) setting default parameters. Tree output was edited using Adobe illustrator (2019).

### SDS-PAGE and Western Blot Analyses

Protein electrophoresis was performed following the method described by Laemmli under reducing conditions (24). Eluted proteins from affinity purification experiments were mixed with 1X Laemmli sample buffer (Bio-Rad), resolved in 4-20% Mini-PROTEAN®TGX precast protein gels (Bio-Rad) and stained with Pierce® silver stain kit, following manufacturer instructions (Thermo). For proteomic analysis, 50 µl from eluted samples were loaded into a pre-cast gel and run until proteins get stack in a 2 cm-gel area. Then, proteins were counterstained with simply blue safe stain (Invitrogen) and gel pieces were excised for further analysis by MS. For western blot analysis, total protein homogenates were prepared from parasites lysed in RIPA buffer (150 mM NaCl, 20 mM Tris-HCl, pH 7.5, 1 mM EDTA, 1% SDS and 0.1% Triton X-100 added of 1X cOmplete EDTA-free protease inhibitor mixture and 1 mM PMSF). Lysates were kept on ice for 30 min and then centrifuged for 12,000 x *g* for 30min at 4°C. Supernatants were used for protein dosage and then 50 µg of each sample were separated by SDS-PAGE. Electrophoresed proteins were transferred to PVDF membranes using a Bio-Rad transblot apparatus. Following transfer, membrane blots were blocked using the Odyssey® blocking solution (Li-Cor, Biosciences) by 1 h, at RT. Blots were probed with anti-c-Myc tag polyclonal antibody (Thermo) diluted (1:2,000) in 5% skimmed milk in PBS containing 0.3% Tween-20 (PBST) and anti-*α*-tubulin monoclonal antibody (Sigma) (1:45,000) for 1 h. After washing three times with PBST, the blots were incubated with IRDye® 800CW goat anti-mouse IgG and IRDye® 680LT goat anti-rabbit IgG (Li-Cor systems) diluted in PBST (1:10,000). Immunopurified proteins were also stained with Pro-Q® Diamond phosphoprotein gel dye (Thermo) or Phospho-Tag (ABP Biosciences, MD, USA) and visualized in a ChemiDoc MP imaging system (Bio-Rad systems) under specific emission/excitation wavelengths (550/580 nm).

### CRISPR/Cas9-mediated Endogenous Tagging

For validation experiments of putative protein substrates of 5-IP_7_, we chose four different candidates identified under both conditions in our screen (Mg^2+^ and EDTA). Then, we deployed a CRISPR/Cas9 gene editing strategy to modify the C-terminus and target proteins were immunoprecipitated by using commercial antibody-coated beads for protein pyrophosphorylation analysis. For that, the Cas9/pTREX-n vector was digested with BamHI to clone a -gene-specific single guide RNA (sgRNAs) targeting the 3’ extremity of five different GOIs: TcAc_CPI (TritrypDB accession number: TcCLB.510857.10), TcBAR (TcCLB510515.30), TcMotorKinesin (TcCLB506211.114), TcIP6K (TcCLB504213.90) and TcCK1 (TcCLB.508541.230), encoding for the hypothetical choline/carnitine *o*-acetyltransferase-domain containing protein, hypothetical BAR-domain containing protein, C-terminal motor-kinesin and inositol hexakisphosphate kinase, respectively specific primers (primers sense 1, 8, 13, 18, and 2 used as antisense, Table S5). Cloning was performed by standard procedures using T4 DNA ligase (Promega) and the procedure was the same as described previously (25). Plasmid DNAs from each one of these constructs (3’-end sgRNA/Cas9/pTREX-n) were prepared in nuclease-free water and co-transfected with a DNA donor cassette containing the antigenic protein tag named spaghetti monster-cMyc tag (26). The hygromycin resistant gene was used as a selectable marker to induce homology-directed repair. Templates for homologous recombination were amplified by PCR with 120-nt ultramers, of which 100-nt correspond to regions located right upstream the stop codon and downstream of the Cas9 3’-cleavage site for each GOI and 20-nt for annealing with consensus regions present in the vector pMoTag-smMyc used as a template (primers 4-5, 9-10, 14-15 and 19-20, Table S5). A TEV cleavage sequence was included in the reverse ultramer. The pMoTag-smMyc construct was generated in our laboratory similarly to that recently described (27). PCRs were amplified using Q5® High fidelity DNA polymerase 2X master mix (NEB), 25 pmol of specific ultramers and 20 ng of DNA template. PCR cycling conditions were: initial denaturation for 2 min at 98°C, followed by 35 cycles of 20 s at 98°C, 10 s at 60°C and 1 min at 72°C followed by a final extension for 5 min at 72°C. Then, 25µg of sgRNA/Cas9/pTREX-n and donor template were co-transfected to epimastigote forms and cultured for 5-weeks in the presence of hygromycin and G418 at 250 µg/ml each. After selection, clonal populations were obtained by limiting dilution in conditioned medium (LIT-20% FBS + Hyg + G418) and then seeded at 0.5 parasites/well in a 96-well plate under humid atmosphere. Genetic validation was performed by PCR using specific primers that anneal into a specific locus of GOI out of the homologous recombination regions (primers 6-7, 11-12, 16-17 and 21-22 Table S5).

### Immunofluorescence Analysis

Wild type and endogenously-tagged epimastigotes were washed with BAG buffer and fixed with 4% paraformaldehyde in PBS for 1 h, at RT. Cells were adhered to poly-lysine-coated coverslips and then permeabilized for 5 min with 0.3% Triton X-100. After permeabilization, cells were blocked with PBS containing 3% BSA (w/v), 1% fish gelatin (v/v), 50 mM NH_4_Cl, and 5% goat serum for 1 h, at RT. Samples were incubated with a monoclonal anti-c-Myc-tag (1:50), diluted in 1% BSA in PBS (pH 8.0), and then incubated for 1 h, at RT. Cells were washed five times with PBS (pH 8.0), and then incubated for 1 h, at RT (light-protected) with Alexa Fluor 488-conjugated goat anti-mouse or Alexa Fluor 546-conjugated goat anti-mouse antibody (1:1,000). Then, samples were washed three times and mounted on slides using Fluoromount-G mounting medium containing 3 µg/ml of 4’,6-diamidino-2-phenylindole (DAPI) to stain DNA. As a control, WT cells were incubated with anti-cMyc. Differential interference contrast and fluorescence optical images were captured on a Delta vision II inverted microscope system (Olympus IX-71) with a 100X objective. Z-series were acquired and deconvoluted using the SoftWorx software.

### Immunoaffinity Purification of Putative 5-IP_7_ Protein Substrates

The protein products from C-terminally tagged epimastigotes were immunopurified for subsequent analysis in the presence of 5-IP_7_. Mutant parasites were grown in selective LIT media (Hyg + G418) at log-phase (≤ 2×10^7^ parasites/ml), as described above. Parasites were harvested (1,600 x *g* for 7 min) and washed twice with BAG. Cell pellets were resuspended in sucrose buffer (150 mM sucrose, 20 mM glutamate-sodium, 20 mM Hepes-KOH, pH 7.7, 3 mM MgCl_2_) and centrifuged at 3,300 x *g* for 10 min at 4°C. Pellets were resuspended in the same buffer and incubated by 20 min on ice. One mM dithiotreitol (DTT) cOmplete EDTA-free protease inhibitor mixture, and 1 mM PMSF were added to the suspension, and mixed gently with a Dounce homogenizer (type B pestle). Aliquots of 1 ml were stored at −80°C for further use. Samples were added to 1/10 of extraction buffer (1.5 M KCl, 20 mM Tris-HCl, pH 7.7, 3 mM MgCl_2_, 0.5 mM DTT, 1% Tween-20) and mixed by inversion. Lysates were clarified by centrifugation for 21,000 x *g* for 15 min at 4°C and supernatants were collected. Proteins were incubated with 75 µl of Pierce™ anti-c-Myc magnetic beads (Thermo), previously equilibrated with binding buffer (150 mM NaCl, 20 mM Tris-HCl, pH 7.7, 3 mM MgCl_2_ and 0.05% Tween-20 buffer), and incubated for 1 h, at 4°C under constant mixing. Beads were collected with a magnetic stand, washed twice with 5X TBS-T buffer (0.75 M NaCl, 125 mM Tris-HCl, 0.25% Tween-20) and finally washed with ultrapure water. Proteins were eluted from beads by adding 50 U of AcTEV™ protease (Thermo) in 50 mM Tris-HCl, pH 8, and 0.5 mM EDTA for 4 h, at 4°C. An additional elution step with 50 mM NaOH buffer was also performed and neutralized with Tris solution. Protein samples were mixed with 4X Laemmli buffer (Bio-Rad) for protein analysis.

### Assessment of Protein Pyrophosphorylation

For analysis of 5-IP_7_-driven protein phosphorylation, radiolabeled 5ß[^32^P]5-IP_7_ was generated by enzymatic synthesis using recombinant mouse IP6K1 in the presence of *γ*[^32^P]-ATP, as described previously (28). Ten reactions containing 1X hot reaction buffer [20 mM Tris-HCl, pH 7.4, 50 mM NaCl, 6 mM MgSO_4_, 0.05 mM ATP-Mg, 1 mM DTT, 5 mM creatine phosphate, 1 U/ml creatine kinase, 200 µCi of *γ*[^32^P]-ATP (Perkin Elmer), 200 µM IP_6_-Na salt and 0.2 µM of His-IP6K1] were incubated for 16 h at 37°C in a total volume of 50 µl. Purification of 5ß[^32^P]5-IP_7_ was performed by precipitation with magnesium and chelation, following a method recently described (29). Reaction products were pooled, and the sample was passed through a short C18 column (SepPak Vac RC tC18 100 mg, Waters) pre-equilibrated with 5 ml of acetonitrile and 5 ml H_2_O. The column was washed with 0.25 ml aliquots of water until no radioactivity remained. MgCl_2_ was added to the filtrate to a final concentration of 16 mM. The sample was precipitated at room temperature and the pH was adjusted to 8.5-9 with 0.1 M NaOH. The sample was then centrifuged for 5 min at 5,000 x *g* and the precipitate was washed three times with 0.25 ml of 8 mM MgCl_2_ (pH 9). The precipitate was resuspended in 0.1 ml of NH_4_HCO_3_ buffer (10 mM, pH 8) and mixed with Chelex 100 resin (0.3 ml wetted bed volume, pre-equilibrated with 10 mM NH_4_HCO_3_, pH 8). The buffer/resin suspension was added to a Polypropylene-reactor 2 ml with polyethylene frit loaded with Chelex 100 column (1 ml bed volume, pre-equilibrated with 10 mM NH_4_HCO_3_, pH 8) for chelation of excess Mg^2+^. The product was flushed through the column with water until no radioactivity remained in the column and the eluent was collected. To the eluent, 10X apyrase buffer (New England Biolabs, MA, USA) and 4 U/ml apyrase (NEB) were added and the reaction was incubated overnight at room temperature. Then, another precipitation and chelation steps were repeated. The final eluate of the Chelex column was collected and concentrated in a speed-vac evaporator to afford the ammonium salt of the product as a concentrated solution. A reaction without adding hot ATP was run in parallel to be analyzed by PAGE. 5-IP_7_-mediated phosphorylation was assayed using parasite whole lysates, protein samples obtained from C-terminally tagged parasites, and recombinant proteins produced in *E. coli*. Cell-free whole lysates were prepared by collecting 2 × 10^8^ parasites and resuspending them in phosphorylation buffer I: 20 mM Hepes, pH 6.8, 1 mM EGTA, 1 mM EDTA, 0.1% (w/v) CHAPS, 5 mM DTT, 5 mM NaF, 1 mM PMSF, and 1X cOmplete EDTA-free protease inhibitor cocktail, for 15 min on ice, as detailed before (28). Lysis was completed by passage through an insulin syringe, the lysate was clarified (20 min at 15,000 x *g* at 4°C) and the supernatant was used for protein dosage. Then, 200 µg of protein lysates were incubated with 10^5^ dpm of 5ß[^32^P]5-IP_7_ at 37°C for 40 min. Briefly, eluted samples from TcAc_CPI-cMyc-, TcBAR-cMyc-, TcC-motorkinesin-cMyc- and TcIP6K-cMyc-tagged cell lines and GST-NSR1 (used as control) were incubated with 1 µCi of 5ß[^32^P]5-IP_7_ in phosphorylation buffer II: 20 mM Tris-HCl, pH 8, 50 mM NaCl, 6 mM MgCl_2_, 1 mM DTT, 0.5 mM NaF for 40 min at 37°C. For assays with recombinant proteins produced from bacteria, proteins were pre-treated with 0.5 mM ATP and 2.5 U casein kinase 2 (CK2) in 1X CK2 buffer (NEB) for 15 min at 30°C. To rule out false positives from traces of hot ATP, a control treatment in the absence of cold ATP was also included. After incubation, 1X LDS sample buffer (Invitrogen) were added and the sample was incubated for 10 min at 70°C. Proteins were resolved in 4-12% NuPAGE® Bis-Tris gels (Invitrogen) for 50 min and 200 V, at 4°C. Then, proteins were blotted to PVDF membranes using standard procedures (1 h x 100 V at 4°C). Membranes were dried out at RT and then exposed to a phosphoscreen cassette (GE) for 6 days for further scanning in a phosphoimager analyzer (Bio-Rad).

### Phosphoprotein Analysis of TcAc_CPI

To analyze the phosphorylation state of TcAc_CPI, we performed mutagenesis studies of the polyserine tract identified as putative site for protein pyrophosphorylation (10). The full TcAc_CPI cds (TcCLB.510857.10) was cloned into the pET32-Ek/LIC vector (Novagen®) resulting in the pET32/TcAc_CPI construct (primers 33, 34, Table S2). We prepared mutants with phosphomimetics (Ser^579^Asp) or full deletion of the polyserine tract (Ser^578-580^) using the pET32/TcAc_CPI construct as template. Specific primers were designed following the instructions of the Q5 site-directed mutagenesis kit (NEB) (primers 35-38, Table S2) and mutations were verified by DNA sequencing (primers 39-40, Table S2). The three protein variants products were expressed in *E. coli* C43 host strain and purified by affinity chromatography, as detailed for TcIP6K experiments. Protein products were *in vitro* phosphorylated with CK2 as described for GST-Nsr1, incubated (37°C for 40 min) with increased concentrations of 5-IP_7_ (0-3µM), and mixed with LDS 1x sample buffer (70°C for 10 min). Protein samples were resolved by NuPAGE gel electrophoresis and stained with Phospho-Tag followed by free stain dye for total protein as indicated by the manufacturer (ABP Biosciences, MD, USA).

## RESULTS

### Presence of IP_6_ and 5-IP_7_ in the Proliferative Stages

Using an LC-MS based analytical detection method, we investigated the presence of IP_6_ and 5-IP_7_ in both proliferative stages of *T. cruzi*, the insect culture form or epimastigote, and the intracellular form or amastigote. We observed that both IP_6_ [M-1H: 659 m/z] and IP_7_ [M-2H: 369 m/z]^2-^ were detected in cell-free extracts from both life cycle stages (Fig. 1A, B). The monoisotopic molecular mass of 5-IP_7_ [M-1H: 739 m/z] was less abundant than its double deprotonated ion precursor [M-2H]^-2^, as seen with the LC-MS profile of 5-IP_7_ used as standard (Fig S1B). The concentration of 5-IP_7_ was calculated as 2.92 ± 0.57 µM (Fig. S1B) when extracts of different amounts of epimastigotes were prepared. Amastigotes possess about twice that amount (Fig. 1B). We then analyzed the potential protein substrates for 5-IP_7_ by combining protein affinity enrichment and bioinformatic analysis as detailed in Fig. S2.

**Figure 1.**
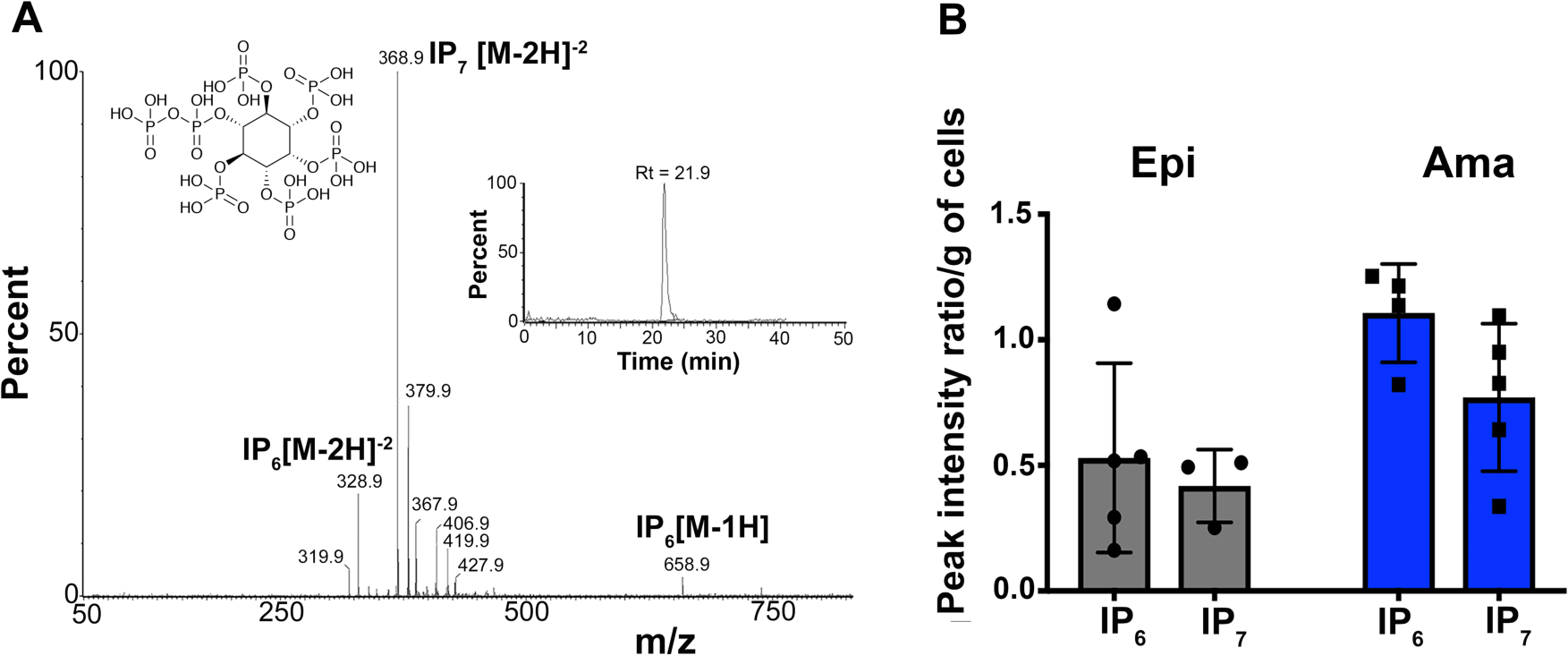
(A) Negative ion electrospray mass spectrometry of inositol hexakisphosphate (IP_6_) and 5-diphosphoinositol-pentakisphosphate (5-IP_7_) separated by liquid chromatography from epimastigote extracts. Inset shows chemical structure of and LC chromatogram for 5-IP_7._ The structure of IP_6_ is shown in Fig. S1. (B) Comparison of IP_6_ and 5-IP_7_ levels in epimastigote and amastigote extracts. Levels are expressed relative to peak integration of internal standard and normalized per mass of wet pellet. Values are means ± s.e.m. of three independent biological experiments (*n* = 3). Differences between values in epimastigotes and amastigotes were significant (p < 0.05, Student’s *t* test).

### An IP_7_-affinity Reagent Identifies Potential Protein Substrates in Proliferative Stages

To identify potential protein substrates that bind to 5-IP_7_, we used affinity-based probes previously developed to detect the 5-IP_7_ interactome of *Saccharomyces cerevisiae* (30). We used resin-bound 5PCP-IP_5_ (a non-hydrolyzable analog of 5-IP_7_) and control reagent C, which contains the same Affi-Gel 15 resin and polyethylene glycol (PEG) linker as 5PCP-IP_5_ but with a negatively charged phosphate head group to detect nonspecific electrostatic interactions and serve as control (Fig. S2). Equal amounts of epimastigote and amastigote lysates were applied to resin-bound 5PCP-IP_5_ and reagent C, followed by washes with buffer and elution with excess IP_6_ (Fig. 2A). The eluted fractions were analyzed by mass spectrometry as described under Experimental Procedures. Total lysates were spiked with human diphosphoinositol polyphosphate phosphohydrolase 1 (HsDIPP1), a protein that interacts with 5-IP_7_ and was used as a positive control in our protein identification assays. Experiments were done in the presence of either 1 mM MgCl_2_ or 1 mM EDTA during affinity enrichment, washing, and elution steps. Three independent biological experiments were carried out and we considered only proteins that were enriched four-fold or higher, as assessed by their relative spectral counts, by 5PCP-IP_5_ compared with reagent C (Fig. S3 and Tables S1-S4).

**Figure 2.**
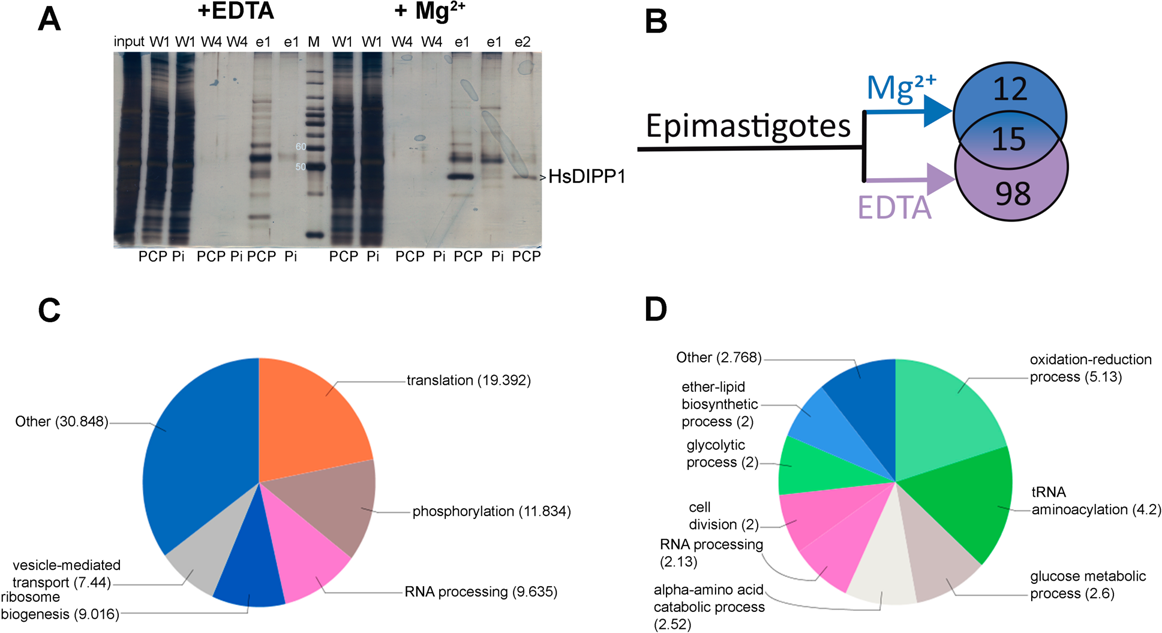
Enrichment of epimastigote proteins by interaction with 5PCP-IP_5_/phosphate affinity reagents. (A) Whole epimastigote lysates (input) were incubated with 5PCP-IP_5_ (PCP) and control reagent C (phosphate, Pi) beads under both metal-free conditions (+EDTA) and magnesium-added (+Mg^2+^) conditions. After successive washing steps (w1 and w4) samples were eluted twice with IP_6_ (e1 and e2) and resolved by SDS-PAGE followed by silver staining. hDiPP1 is the standard marker. MW markers (M) are between samples. (B) Number of proteins isolated under both conditions in the epimastigote stage. (C) Functional classification of proteins identified in the 5PCP-IP_5_-binding proteome of epimastigotes under metal-free conditions. (D) Functional classification of proteins identified in the 5PCP-IP_5_-binding proteome of epimastigotes in the presence of magnesium ions.

Our experiments with 5PCP-IP_5_ reagent in the absence of magnesium (metal-ion depleted conditions) isolated about 120 proteins from each stage (Fig. 2B, 3A). Proteins involved in translation processes, phosphorylation, ribosomal biogenesis, RNA-processing elements, and vesicle-mediated intracellular transport were enriched in epimastigotes (Fig. 2C). In the amastigote stage, there was an overrepresentation of similar biological processes as seen in epimastigotes but also elements involved in intracellular transport, response to stress and oxidation-reduction processes were isolated (Fig. 3B). When magnesium was present, we identified a smaller cohort of proteins (32 hits) in both stages (Fig. 2D and 3C). In epimastigote forms, proteins involved in glycolysis, lipid metabolism, tRNA aminoacylation, cell division, and RNA processing were enriched (Fig. 2D). In amastigotes, we identified proteins involved in translation, intracellular protein transport, maturation of rRNA, protein phosphorylation, cell redox homeostasis, actin nucleation and nucleosome assembly (Fig. 3C). The cellular distribution of these proteins also presented differences between both stages. Nuclear, cytoplasmic and vacuolar proteins were identified in amastigotes and glycosomal, nuclear, membrane-bound and cytoplasmic proteins were enriched in epimastigotes (Fig. 3D, Table S1, S3). There was only a 28% overlap when protein enrichment between epimastigote and amastigote forms was compared under both conditions and the most overrepresented GO-terms were phosphate ion binding (P-value: 1.28e^-10^) and snoRNA binding (P-value: 7.58e^-5^) (Tables S2, S4). In addition, the host vimentin (P84198), actin, cytoplasmic 1 (Q76N69) and the cadherin-related family member 2 (Q9BYE9) were identified in amastigotes devoid of divalent cations and the peroxiredoxin-2 (P32119) was isolated when magnesium was present. Only the actin, cytoplasmic 1, was present under both conditions (Fig. 3A).

**Figure 3.**
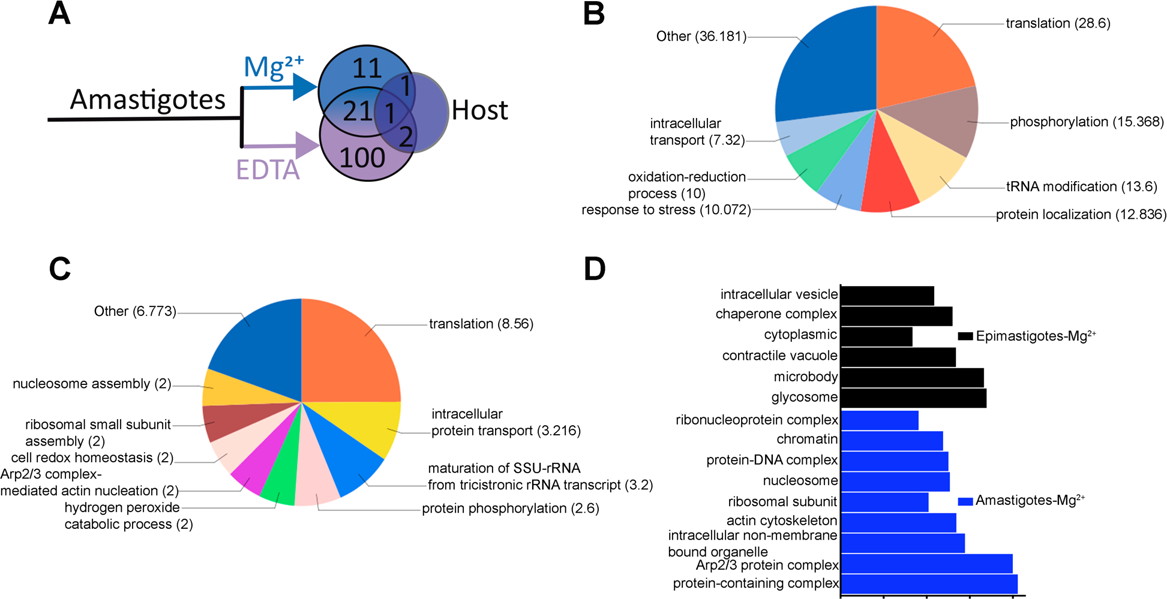
(A) Number of proteins isolated under both conditions in the amastigote stage. (B) Functional classification of proteins identified in the 5PCP-IP_5_-binding proteome of amastigotes under metal-free conditions. (C) Functional classification of proteins identified in the 5PCP-IP_5_-binding proteome of amastigotes in the presence of magnesium ions. (D) Comparison of GO-terms (based on cellular compartment) found in proteins from both stages enriched under the presence of magnesium.

Among the most important candidates, we detected previously- and newly-identified protein targets. Our control protein HsDIPP1 was isolated in the absence of magnesium (Fig. 2A). We were also able to isolate proteins whose homologs have been previously reported as targets for 5-IP_7_. Two proteins that are known to contain an SPX-domain at its N-terminus, were isolated in our screen in the presence of magnesium (Table S2). The phosphate sodium symporter Pho91 (annotated as sodium/sulfate transporter: TcCLB.511671.149) and the catalytic component of vacuolar transporter chaperone complex (VTC) Vtc4 (TcCLB.511127.100/TcCLB.511249.44) were also isolated when magnesium was present in amastigote lysates. A casein kinase type I (TcCLB.508541.225) and a member belonging to the family of calcineurin-like phosphoesterases annotated as kinetoplastid-specific phosphoprotein-phosphatase (TcCLB.504411.30) was isolated in the presence of magnesium in amastigotes (Table S4). Interestingly, this protein has homology to the *S. cerevisiae* endopolyphosphatase Ppn2, which localizes to the acidocalcisome-like vacuole (31).

In order to rule out any non-specific enrichment of highly-abundant proteins in the parasite, data were compared with those reported in the *T. cruzi* proteome of both epimastigotes and amastigotes (32). We only observed a correlation for TcCLB.506211.114, TcCLB.506247.220, TcCLB.511573.58, TcCLB.506211.160 and TcCLB.509505.10, which have been reported as abundant in both proliferative stages, and were isolated under metal-free conditions. TcCLB.506211.114 was also isolated in the presence of magnesium.

### 5-IP7 as Possible Mediator of Vesicle Trafficking and Membrane Dynamics

Our screen identified protein components that participate in the three major steps (assembly, membrane curvature, and fission) of the clathrin-mediated endocytic uptake of proteins and lipids. A clathrin coat assembly protein (TcCLB.509875.190 or TcAP180), was isolated in epimastigotes under cation restriction. TcAP180 localizes to the bladder of the contractile vacuole complex (33). A highly abundant motor kinesin protein (TcCLB.506211.114) was enriched in both stages under both conditions. This protein contains both an epsin N-terminal homology (ENTH) and a P-loop nucleoside triphosphate hydrolases domain. Two hypothetical proteins (TcCLB.507907.50 and TcCLB.511421.10) containing a pleckstrin-homology (PH) domain were isolated from epimastigotes. Also, the Sec3, Sec6, and Sec10 proteins that are part of the exocyst complex were enriched in epimastigotes under cation restriction (TcCLB.510173.10, TcCLB.506367.120, and TcCLB.503829.70, respectively). A hypothetical protein grouped into the Bin/Amphiphysin/Rbs (BAR) domain superfamily (TcCLB.510515.30) was also identified and there was another one containing the same protein domain (TcCLB503965.39). The GTPase annotated as a Rho GTPase-binding/formin homology 2 and −3 (TcCLB.511393.30) and the GTPase activating protein annotated as disgorgin (TcCLB.508723.80) were also identified in epimastigotes (Tables S1 and S3). Interestingly, disgorgin is also a contractile vacuole protein (33).

### 5-IP7 as a Regulator of Purine and Pyrimidine Biosynthetic Steps

Enzymes from the purine and pyrimidine biosynthetic pathways were also enriched in our screen, such as the uridine 5-monophosphate synthase (TcUMPs) (TcCLB.507059.60), which is characterized for being a bifunctional enzyme in trypanosomatids (34) comprising orotate phosphoribosyl transferase (OPRT) and orotidine monophosphate decarboxylase (OMPDC) and for catalyzing the synthesis of UMP, a precursor for pyrimidine nucleotides. We also observed an overrepresentation of proteins acting on nucleotide and purine biosynthesis (35). The enzymes ribose-phosphate pyrophosphokinase [EC 2.7.6.1], guanosine 5-monophosphate reductase (GMP-reductase) [EC 1.7.1.7] and inosine-5-monophosphate dehydrogenase (IMPDH) [EC 1.1.1.205] were enriched in amastigote forms under metal-free conditions (Table S3). These enzymes participate in the nucleotide and purine salvage biosynthetic pathways, respectively. The GMP-reductase (TcCLB.506519.130) catalyses the irreversible NADPH-dependent deamination of GMP to IMP. Interestingly, our study also revealed that IMPDH may interact with 5-IP_7_. TcIMPDH catalyzes the reversible conversion of inosine-5-monophosphate into xanthosine 5-monophosphate which is needed for guanosine biosynthesis. Another enriched protein was the ribose-phosphate pyrophosphokinase, also known as phosphoribosyl pyrophosphate transferase, which participates in the production of PRPP (phosphoribosyl pyrophosphate) from ribose-5-phosphate (R5P), which is needed as a building block for nucleotide synthesis. Interestingly, we found an SPX-domain present at the N-terminus of this enzyme, which is a unique feature conserved across species of the order Kinetoplastida. SPX-domains mediate binding of proteins to inositol polyphosphates (11).

### 5-IP7 and its Putative Role in Acetyl-CoA Metabolism

We also found an overrepresentation of proteins participating in lipid and branched chain amino acids metabolism. The 3-ketoacyl-CoA thiolase (TcCLB.510507.20), found in epimastigotes under metal restriction, is a CoA acetyltransferase that condenses two units of acetyl-CoA producing acetoacetyl-CoA, which is an intermediate of the mevalonate pathway. Under the same conditions, another enzyme of the acetyl-CoA metabolism that was enriched was the acyl-CoA dehydrogenase (TcCLB.509153.120). This is a FAD-dependent mitochondrial membrane protein, that is important because it is capable of transferring electrons to the ubiquinone pool in the mitochondria (36). In amastigotes, under metal restriction, we found an enzyme (annotated as trifunctional enzyme alpha subunit: TcCLB.509701.10) that may be acting on acetoacetyl-CoA reduction resulting in hydroxybutanoyl-CoA. On the other hand, in the presence of magnesium ions we enriched the 3-methylcrotonoyl-CoA carboxylase ß-subunit (TcCLB.506773.40) and a protein containing a choline/carnitine_acyl transferase domain annotated as hypothetical (TcCLB.510857.10). The former is involved in the leucine degradation pathway and the latter may have an activity in the transference of CoA from carnitine that is needed for lipid catabolism (Tables S1, and S3).

Only about ten percent of the proteins identified in cells devoid of divalent ions have been previously identified as phosphoproteins by mass spectrometry studies (37). Six hypothetical proteins (TcCLB.506925.110, TcCLB.510515.30, TcCLB.510409.50, TcCLB.506945.320, TcCLB.511181.140, TcCLB.509011.60), the clathrin coat assembly protein (TcCLB.503449.30) and C-terminal motor kinesin (TcCLB.506211.114) were found in epimastigotes datasets and pyruvate phosphate dikinase (TcCLB.510101.140), beta tubulin (TcCLB.506563.40), clathrin coat assembly protein (TcCLB.509875.190), C-terminal motor kinesin (TcCLB.506211.114) and two proteins annotated as hypothetical (TcCLB.509859.10 and TcCLB.511181.140) were found in amastigotes. Under the presence of magnesium, the succinate dehydrogenase subunit (TcCLB.511643.10), I/6 autoantigen, putative protein (TcCLB.508323.100) and the C-terminal kinesin (TcCLB.506211.114) were found in amastigotes whereas clathrin coat assembly protein (TcCLB.503449.30), C-terminal motor kinesin (TcCLB.506211.114) and the hypothetical (TcCLB.510515.30) have phosphoproteome evidence in epimastigotes.

### Validation of 5-IP_7_ Protein Substrates

To further corroborate that the protein substrates found in our screen interact with 5-IP_7_, we designed a CRISPR/Cas9 strategy to endogenously tag four candidates (TcAc_CPI, TcCK1, TcBAR, and TcMotK) at their C-terminus using the antigenic spaghetti monster tag (sm-cMyc) (26) preceded by a TEV protease cleavage site in epimastigotes (Fig. 4A). We selected candidates in order to achieve a diversity of structural motifs and potential cellular localization. The TcAc_CPI (TcCLB.510857.10) and TcCK1 (TcCLB.508541.230) were enriched in the presence of Mg^2+^, while TcBAR (TcCLB.510515.30) and TcMotK (TcCLB.506211.114) were enriched under metal-free conditions. TcCK1 and TcMotK were obtained from amastigote lysates, while TcBAR and TcAc_CPI were obtained from epimastigote lysates. The IP6K of *T. cruzi* (TcCLB504213.90, TcIP6K) was used as negative control since it naturally binds to IP_6_. Mutant cell lines in both alleles were obtained for the TcBAR, TcIP6K and TcAc_CPI genes (Fig. 4B). Genetic validation using specific primers showed the insertion of the sm-cMyc cassette and selection marker in the right locus (amplicon size ≈3 kb). Mutants for C-terminal motor kinesin (MotK) gene resulted in a heterozygous clone as both, the endogenous and mutant copies of the gene, were detected (Fig. 4B). Protein detection of these products using commercial antibodies (anti-c-Myc) showed molecular weights compatibles with the expected ones. The fusion of sm-cMyc tag to the target protein produces a 37 kDa increase in the molecular size. Then, corresponding molecular sizes observed were: 110, 140, 165, 150 kDa for TcBAR, TcAc_CPI, TcMotK, and TcIP6K kDa, respectively (Fig. 4C). Anti c-Myc labelling showed no cross-reaction with wild type cells. The presence of other protein bands suggests proteolysis events that were only evidenced in the mutant cell lines, as they harbour a TEV protease cleavage site. The TcBAR tagged cells displayed labelling around the plasma membrane and within intracellular vesicles near the flagellar pocket (Fig. 4D). TcAc_CPI, TcIP6K and TcMotK seem to be cytosolic proteins as their labeling extends throughout the flagellum, although TcAc_CPI might be present in the mitochondria (Fig. 4D). TcCK1 was detected using anti-cMyc in IFA assays but not by immunoblotting (data not shown). Tagged proteins TcBAR, TcAc_CPI, TcMotK, and TcIP6K were successfully immunopurified from total-protein lysates of epimastigotes. To evaluate whether the selected parasite candidates are phosphoproteins, they were submitted to in-gel detection using ProQ diamond dye; a commercial reagent selective to stain phosphoproteins (38). TcAc_CPI (Fig. 5A) and TcBAR (Fig. S4C) were labelled with ProQ diamond.

**Figure 4.**
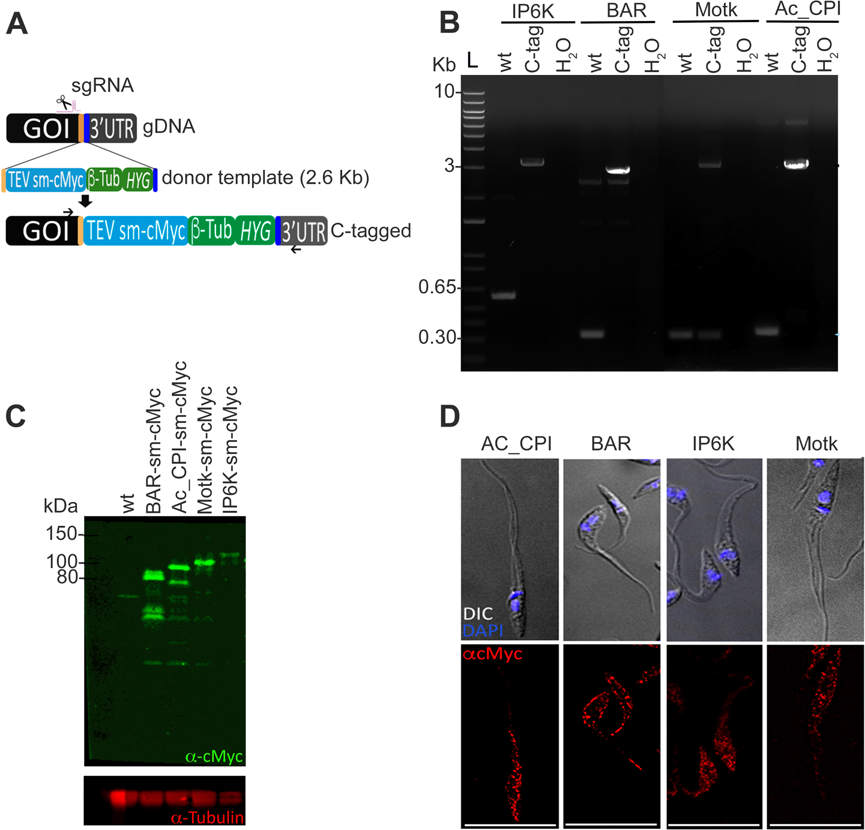
C-terminal tagging of 5PCP-IP_5_-binding proteins. (A) Schematic representation of CRISPR/Cas9-mediated endogenous tagging. Gene-specific sgRNAs were used to target the insertion of a donor template (2.6 Kb) flanked by homology arms (100 nt length), and harbouring a TEV-protease cleavage site fused to sm-cMyc sequence to be expressed in frame with genes of interest (GOI). The intergenic region of β-tubulin (*β-tub*) gene drives the expression of the hygromycin (*HYG*) resistance gene, used as selection marker. Specific primers were designed for validation of cell lines (*arrows*). (B) Verification of endogenous tagging by PCR analysis. Lanes on 1% agarose gel: L, ladder; and PCR products amplified using gDNAs from parental (wt), resultant clonal populations (C-tag) and negative control (H_2_O). A fragment of approximately 300-600 bp is amplified in wt cells, which corresponds to the 3’-end of each GOI and its 3’-UTR. After transfection and selection of mutants, the C-tagged cell lines contain a larger fragment (∼3 Kb). The GOI used for tagging were: inositol hexakisphosphate kinase (IP6K), hypothetical BAR domain-containing protein (BAR), motor kinesin (MotK), and hypothetical choline/carnitine *o*-acetyltransferase-domain-containing protein (Ac_CPI). Endogenous tagging was verified by western blot (C) and immunofluorescence analyses (IFA, D). The sm-cMyc tag adds a fusion product of 37 kDa to the protein of interest. Mouse monoclonal antibody anti-cMyc was used for protein detection and mouse anti-alpha-tubulin was used as loading control. (D) Immunofluorescence assays of tagged epimastigotes using anti-cMyc. Transfected parasites were incubated with mouse anti-cMyc and stained with mouse anti-IgG conjugated to Alexa fluor-546 secondary antibody. DAPI was used for DNA staining and is shown merged to differential interference contrast (DIC) image. Scale bars = 15 µm.

**Figure 5.**
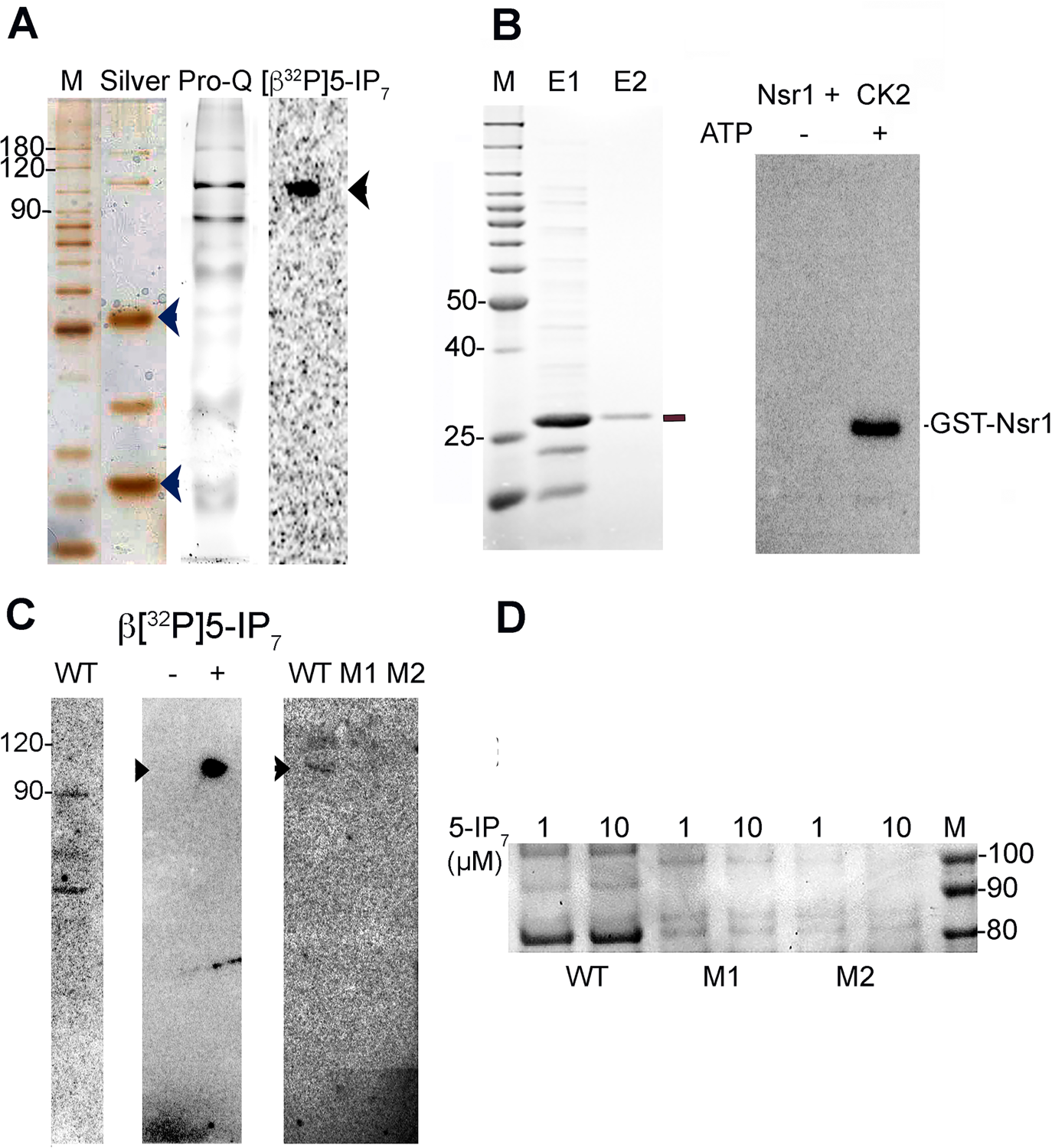
Detection of protein pyrophosphorylation. (A) C-tagged TcAc_CPI was immunopurified from parasite lysates using anti-cMyc-magnetic dynabeads, and phosphorylation and pyrophosphorylation events were analyzed. Silver staining (Silver) of elution products revealed the cMyc-tagged and untagged proteins after TEV cleavage. The heavy (50 kDa) and light chain (25 kDa) of anti-IgG were also co-eluted (*arrowheads*). Eluate was also stained with ProQ dye, which is specific for phosphoprotein detection. After incubation with [β^32^P]5-IP_7_, a positive signal was detected for TcAc_CPI (*arrow*). (B) The GST-Nsr1 (27-50 amino acids) fragment was expressed and purified from bacteria to be used as control in *in vitro* pyrophosphorylation experiments (*left*). [β^32^P]5-IP_7_-driven phosphorylation was observed only after priming with CK2 and ATP (*right*). No traces of hot ATP were seen after precipitation, chelation and apyrase-treatment of radiolabeled 5-IP_7_. (C) 5-IP_7-_driven protein pyrophosphorylation. [β^32^P]5-IP_7_ was incubated with cell-free lysates from epimastigotes (WT, *left*), immunopurified TcAc_CPI-sm-cMyc (*center*) and the mutant TcAc_CPI proteins (M1, S579A; and M2, S^578-80del^) primed with casein kinase 2 (CK2) and ATP prior to the pyrophosphorylation assay. A control without [β^32^P]5-IP_7_ was also included (*center*) (D) Phosphorylation state of TcAc_CPI and variant isoforms produced in bacteria. Recombinant proteins were analyzed with two different concentrations of 5-IP_7_ (1, 10µM), resolved by NuPAGE gels and stained with PhosphoTag. Only the WT version presented positive staining for phosphoproteins.

### TcAc_CPI Undergoes Pyrophosphorylation Events

Of the four candidates analysed, only TcAc_CPI and TcBAR were found to be phosphoproteins, and were isolated under magnesium and metal-free conditions, respectively, so we explored whether they were susceptible to pyrophosphorylation in vitro by ß-phosphoryl transfer from 5-IP_7_ into a phosphoserine residue. Radio-labelled 5-IP_7_ was produced enzymatically using the recombinant mouse-IP6K1 and *γ*^32^P-ATP (Fig. S4A). The product *β*[^32^P]5-IP_7_ was treated with apyrase, precipitated with magnesium, chelated, and eluted for its use in binding experiments with the selected protein targets immunopurified from parasite whole lysates to ensure the necessary pre-phosphorylation by endogenous protein kinases. We also expressed a GST-Nsr1 protein fragment (amino acids 27-50) (Fig. 5B, left panel), which is susceptible to pyrophosphorylation in vitro (10), as positive control. The recombinant GST-Nsr1 fragment was pre-treated with CK2 and ATP before the incubation with *β*[^32^P]5-IP_7_ (Fig. 5B, right panel). Then proteins TcBAR and TcAc_CPI (Fig S4B, tagged and untagged eluates) were immunopurified and treated with *β*[^32^P]5-IP_7_ for 40 min at 37°C. The reaction mixtures were incubated at 70°C for 10 min in sample buffer, resolved by NuPAGE Bis-tris gels, transferred to PVDF membranes, and the ^32^P signals were detected by phosphoimager analysis. Strong signals were observed for TcAC_CPI (Fig. 5A, right) and GST-Nsr1 (Fig. 5B, right) consistent with protein pyrophosphorylation, whereas TcBAR was not modified with the radiolabel (Fig. S4C, right). Absence of residual hot ATP in our radioactive preparation was corroborated by non-phosphorylation of GST-Nsr1 when the ß[^32^P]5-IP_7_ product was treated with CK2 under nominal absence of cold ATP (Fig. 5B, right panel).

Incubation of epimastigote cell-free lysates with *β*[^32^P]5-IP_7_ detected several radiolabeled proteins showing that other proteins can be susceptible to this post-translational modification (Fig. 5C, left panel). We also purified the recombinant TcAc–CPI from bacteria, as confirmed by antibody detection (Fig. S4B), and its phosphorylation state was analyzed after its treatment with casein kinase 2 (CK2) and ATP (Fig 5C, middle panel). Analysis of the protein sequence of TcAc_CPI revealed the presence of a serine-enriched motif (Ser^578-580^) flanked by acidic residues (Asp^577^ and Glu^583^), which is characteristic of this posttranslational modification (10). To confirm that protein pyrophosphorylation of TcAc_CPI involves serine residues, site-directed mutagenesis on this motif was done and the recombinant 6xHis-TcAc_CPI was isolated. Both a mutant in which the polyserine tract was deleted (Ser^578-580 del^) and a mutant in which serine^579^ was replaced by a phosphomimetic aspartate residue (S579A) were obtained, purified, treated with CK2, and incubated with *β*[^32^P]5-IP_7_. No pyrophosphorylation was observed when Ser^578-580^ tract was removed (Fig. 5C, right panel, M1) or when the serine was replaced by aspartate (Fig. 5C, right panel, M2). When these recombinant forms of TcAc_CPI were stained with Phospho-Tag, only the WT displayed positive staining using this dye (Fig. 5D). These data suggest the Ser^578-580^ tract is the site of phosphorylation in TcAc_CPI. These data also confirm the presence of a phosphoserine residue in TcAc_CPI needed for *β*-phosphoryl transfer from 5-IP_7_.

### TcPRPS4 is a Unique SPX Domain-containing Enzyme Involved in Nucleotide Synthesis

TcPRPS4, which is involved in nucleotide biosynthesis (Fig. S5A), contains a domain characteristic of proteins that binds to 5-IP_7_. Genome data revealed three other different genes encoding for putative phosphoribosyl pyrophosphate synthetases (TcCLB.506321.310, TcCLB.510431.250, and TcCLB.508717.30), which only share 33% sequence identity. The major differences are at the N-terminus, where an SPX domain is present, and at the C-terminus, where an 85 amino acids region is present (Fig. S5B). This stretch has no homology with PRPSs from any species. We named this protein as TcPRPS4 (Fig S5B). This enzyme is unique to the order Kinetoplastida with no homologues in humans or plants (Fig. S6). Its putative activity is the transference of a pyrophosphate from ATP into C-1 of ribose-5-phosphate forming PRPP and AMP (Fig. S5A).

## DISCUSSION

Our studies revealed a number of 5-IP_7_-interacting proteins in the proliferative stages of *Trypanosoma cruzi*. The combined purification by binding to an 5-IP_7_ affinity reagent, with mass spectrometry identification, and bioinformatic analysis, revealed potential 5-IP_7_-regulated processes in epimastigotes and amastigotes of *T. cruzi*. Two groups of putative targets were identified either in the presence of magnesium ions or under metal ion restriction. In the absence of magnesium, there was an overrepresentation of proteins involved in purine metabolism, protein phosphorylation, nucleosome assembly, cell redox homeostasis, parasite-surface components, and exocytosis, and only 37% overlap in both stages. In the presence of magnesium, there was an overrepresentation of proteins involved in tRNA amino acylation, phosphate homeostasis, lipid and glucose catabolism, and cell division, and 15% overlap between both stages. We identified a choline/*o*-acetyltransferase domain-containing phosphoprotein that undergoes 5-IP7-mediated phosphorylation events at a polyserine tract (Ser^578-580^). We also identified a novel SPX domain-containing phosphoribosyl transferase (TcPRPS4).

Inositol pyrophosphates (PP-IPs) have been postulated as signaling molecules because their interaction with proteins regulates cellular homeostasis. The biological importance of 5-IP_7_ is based on the presence and position of its high-energy diphosphate bond. Most of the studies describing the functions of PP-IP_s_ are based in effects observed after downregulating levels of these metabolites as a consequence of deletion of *IP6K1*/*IP6K2*/*PPIP5K* genes. Then, a plethora of cellular activities in yeast and human cells have been uncovered embracing vesicular trafficking (39), chemotaxis (40), insulin secretion(41), autophagy (42), and phosphate homeostasis (16), among others.

The combined use of affinity reagents (5PCP-IP_5_- and C-reagent) allowed the separation 5-IP_7_-protein targets from spurious interactions and revealed 5-IP_7_-protein associations that are ubiquitously distributed at different parasite compartments. We identified substrates containing IP_s_/PP-IP_s_ binding-domains such as Pleckstrin-Homology (PH), Bin/Amphiphysin/Rbs (BAR), VPS-27/Hrs/STAM (VHS) - AP180 N-terminal homology (ANTH), and the most characteristic one the SYG1/Pho81/XPR1 (SPX). 5-IP_7_ protein binding did not need divalent cations, but when magnesium was added its binding targets changed revealing a different set of proteins, as previously reported in yeast studies (22). Magnesium is essential for proteins undergoing 5-IP_7_-driven phosphorylation, which is conserved in this parasite.

We found no significant differences in the relative abundance of IP_6_ when compared with that of 5-IP_7_ in both parasite stages. The 5-IP_7_ levels determined (2.9 ± 0.4 µM) for epimastigote forms are similar to those measured in embryonic fibroblast cells (NIH 3T3), but 20-fold lower than those reported for the amoeba *Dictyostelium* (5-250 µM) (7). In most organisms, levels of IP_6_ are considerably higher than those of 5-IP_7_, due to the lower affinity (mM range) of IP6K for ATP (43). Although kinetic properties of TcIP6K for ATP are unknown, the turnover of ATP varies throughout the proliferative cycle of trypanosomes (3, 0.6 mM in log and stationary phases of growth for epimastigotes, respectively) and relies on parasite’s nutritional status (44).

When protein binding in the absence of magnesium was examined there was an overrepresentation of proteins with annotated phosphate ion binding (GO term: 0042301, P-value 2.91^-10^) and nucleic acid binding (GO term 0003676: 1.68^-4^) terms. One of the most abundant proteins enriched in both stages was a hypothetical protein (TcCLB.511439.40) annotated as phosphate ion binding protein. The protein possesses a putative PH-domain, as predicted by the HHPred server, and in a previous study was identified to bind to polyP (45). The function of this protein is unknown, but the lack of orthologs in other eukaryotes, makes it an interesting candidate for further investigation of its role in phosphate homeostasis in kinetoplastids. *T. cruzi* possesses SPX-domains in the catalytic subunit of the vacuolar transporter chaperone (VTC) complex, which is involved in polyP synthesis and translocation to acidocalcisomes (46), and in the sodium/phosphate symporter (Pho91), involved in Pi release from the contractile vacuole (47). Both proteins (TcCLB.511127.100/TcCLB.511249.44 and TcCLB.511671.149) were detected in amastigote samples.

The role of an actin-related protein 2/3 complex in polyP synthesis has been reported (48). Our study isolated a candidate annotated with this function (TcCLB.504215.40). The protein as well as that encoded by a paralog (TcCLB.506865.10), was also pull down with polyP (45). The protein encoded by TcCLB.504215.40 contains WD-40 repeats (WD40/YVTN) that might be involved in its interaction with inositol phosphate metabolism, as it has been previously shown in a plant inositol polyphosphate 5-phosphatase (49). Interestingly, several other proteins pull down by 5-IP_7_ were also pull down by polyP from epimastigote lysates (6-phospho-1-fructokinase [TcCLB.508153.340], fructose-1,6-bisphosphatase [TcCLB.506649.70], nucleolar protein [TcCLB511573.58], histidine ammonia-lyase [TcCLB.506247.220], serine carboxypeptidase S28 [TcCLB.506425.10] and ribosomal protein L38 [TcCLB.503575.34]) (45).

Previous work studying the action of the IP6Ks demonstrated a role for inositol phosphates in vesicular dynamics and vacuole biogenesis (39, 50). In our screen, we isolated different components whose molecular function is linked to vesicle transport and membrane trafficking in this parasite, like the clathrin coat assembly protein (TcCLB.509875.190 or TcAP180) that localizes to the bladder of the contractile vacuole in epimastigotes (33) and the GTPase activating protein disgorgin (TcCLB.508723.80), also localized to the contractile vacuole complex in *D*. *discoideum* and *T. cruzi*, as seen in proteomic studies (33, 51). In addition to its role in osmoregulation, the contractile vacuole complex of *T. cruzi* is a trafficking hub that is involved in the transfer of GPI-anchored proteins to the plasma membrane (52) and is also a specialized endosomal system that can be used to deliver membrane proteins that are important for the biogenesis of acidocalcisomes (53).

Sec3, Sec6, and Sec 10 proteins, components of the exocyst complex, were also enriched in our screen (TcCLB.510173.10, TcCLB.506367.120, and TcCLB.503829.70) as were BAR-domain containing proteins and GTPases. BAR domains interact with membranes and define curvatures with diverse functions in organelle biogenesis, membrane remodelling, and cell division (54). A protein (TcCLB503965.39) isolated from epimastigotes has a BAR/IMD domain.

A set of several nuclear and nucleolar proteins were identified in our protein screen. The CyaB like adenylyl cyclase, thiamine triphosphatase (CYTH) domain-containing enzyme was found in both epimastigotes and amastigotes (TcCLB.506503.160). We have observed an enrichment of ribosome biogenesis, translational machinery components, and snoRNA binding proteins in our 5-IP_7_-proteome. Amongst the most interesting candidates are a Broad-Complex, Tramtrack and Bric a Brac/POxvirus and Zinc finger (BTB/POZ) domain-containing protein (TcCLB.511277.320) and a protein annotated as Lupus La protein (TcCLB.511367.60). La protein has a canonical role as an RNA polymerase-III transcription factor in the nucleus, but it can also play a role as a translation factor in the cytoplasm.

Few nucleolar RNA-binding proteins with putative functions in pre-mRNA splicing and ribosome biogenesis were found in both stages (TcCLB.508277.230, TcCLB.503773.50, TcCLB.506189.10). There was also enrichment in purine nucleotide biosynthetic enzymes (10 proteins). Similarly, our study identified five different enzymes that participate of purine biosynthetic pathways: PRPS4 (TcCLB.507809.30), IMPDH (TcCLB.507211.40), GMP reductase (TcCLB.506519.130) in amastigotes and PRPS4, UMPS (TcCLB.508373.29) and adenosine phosphoribosyltransferase (TcCLB.508207.74), but no similarities were observed in their protein counterparts compared to the yeast study.

In conclusion, our results suggest the involvement of 5-IP_7_ in a number of metabolic and signaling pathways of the proliferative stages of *T. cruzi*. Some of these pathways, such as ribosome biogenesis, and the regulation of the energy status of the cells have also been reported as 5-IP_7_-dependent in previous studies in yeast (22), while others appear as more specific such as the 5-IP_7_ involvement in the trafficking role of the contractile vacuole of these parasites. Their study could identify new biological functions and reveal novel targets for the chemotherapy of these parasitic disease.

## Acknowledgments

We thank Dennis Philips, head of the Proteomic and Mass Spectrometry (PAMS) Core Facility at UGA, for technical support in LC-MS and proteomic analysis, Robert Puschmann for support in purification of 5ß[^32^P]5-IP_7_, Belen Cassera for assistance in LC-MS analysis, Guozhong Huang for providing the pMoTag-sm-cMyc construct used for tagging, Rodrigo Baptista for assistance in use Blast2Go software, Melissa Storey for technical assistance with amastigote isolation, Henning J. Jenssen for 5-IP_7_, and Muthugapatti Kandasamy and the Biomedical Microscopy Core of the University of Georgia for the use of microscopes.

Funding for his work was provided by the U.S. National Institutes of Health (grant AI107663 to RD).

## Author contributions

B.S.M., N.B., D.F. and R.D. analysed data; B.S.M. and N.B. performed research; B.S.M. and R.D. designed research; B.S.M. and R.D. wrote the paper with contributions of N.B., and D.F.

---

**Figure S1.**
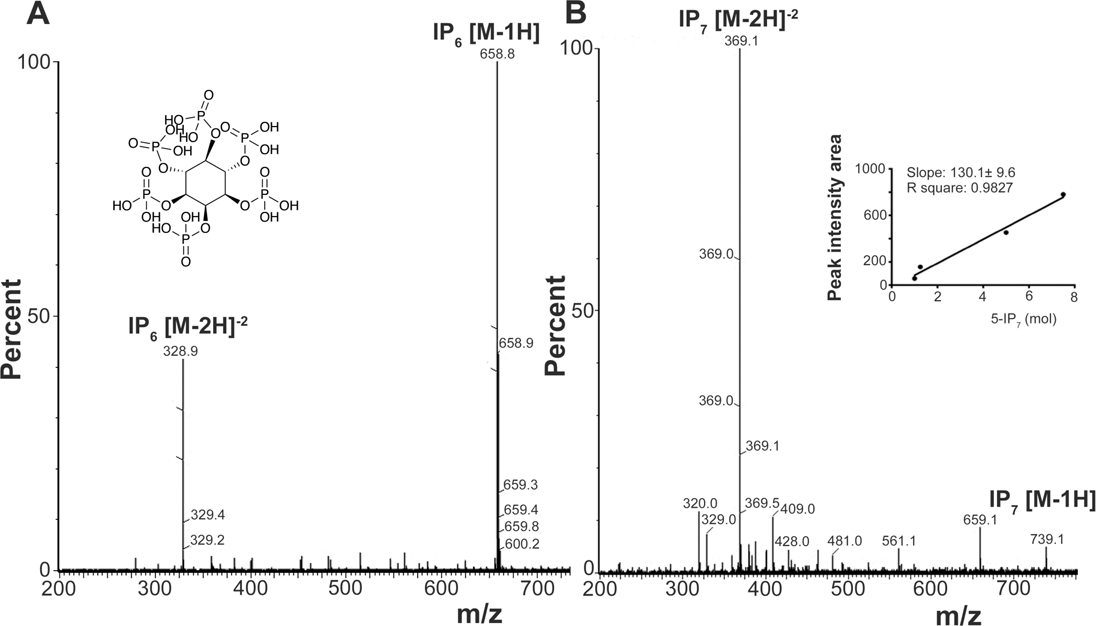
Negative ion electrospray mass spectrometry of inositol hexakisphosphate (IP_6_) (A) and 5-diphosphoinositol-pentakisphosphate (5-IP_7_) (B) standards. (A) Ten nmol of IP_6_ [M = 660 Da] were injected and separated by LC. Two main ions of 329 m/z and 659 m/z (single deprotonated) are shown. Inset shows IP_6_ structure. (B) Ten nmol 5-IP_7_ [M = 740 Da] were injected and separated by LC. Two main ions of 369 m/z (double deprotonated) and 739 m/z (single deprotonated) are shown. Inset shows a calibration curve using 1-10 nmol of 5-IP_7_. Peak integration area was used to determine total amount of 5-IP_7_ in parasite extracts.

**Figure S2.**
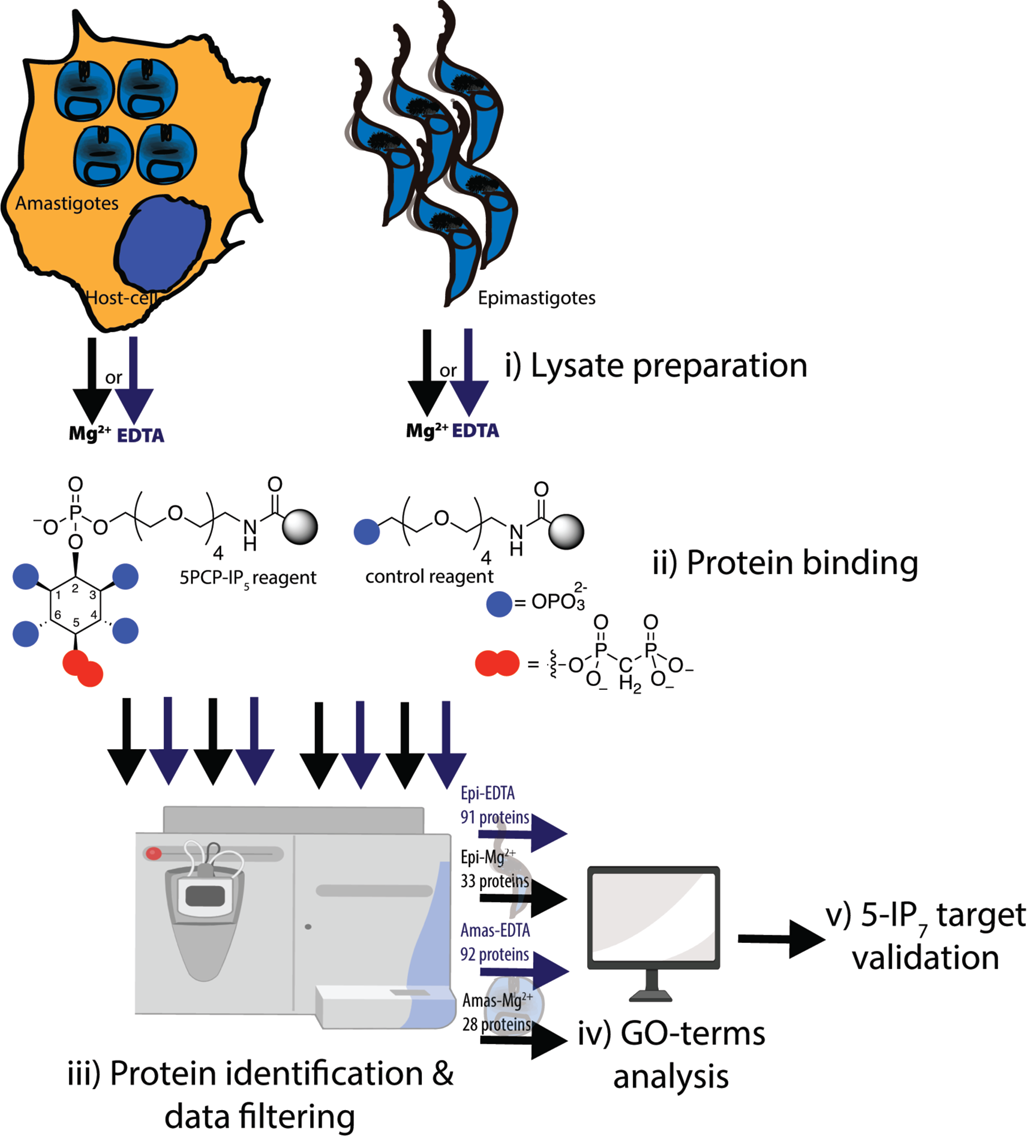
Experimental workflow followed in this study. Cell-free protein samples were prepared from proliferative stages of *T. cruzi*. Epimastigotes were grown axenically and amastigotes were obtained from gamma-irradiated Vero cells infected with cell-derived trypomastigotes. i) Lysates from both stages were prepared in the presence of magnesium ions or under the metal-free conditions. ii) Protein lysates were incubated with 5PCP-IP_5_ or control reagent, separately. iii) Eluted proteins (from 8 different conditions) were identified by mass spectrometry and results were filtered out according to the PSM enrichment obtained in the 5PCP-IP_5_ beads relative to phosphate beads used as control. iv) GO-terms analysis was done for all the resultant dataset hits enriched in at least two biological replicates. v) Based on protein identification data some targets were chosen for validation experiments in the parasite or *in vitro*.

**Figure S3.**
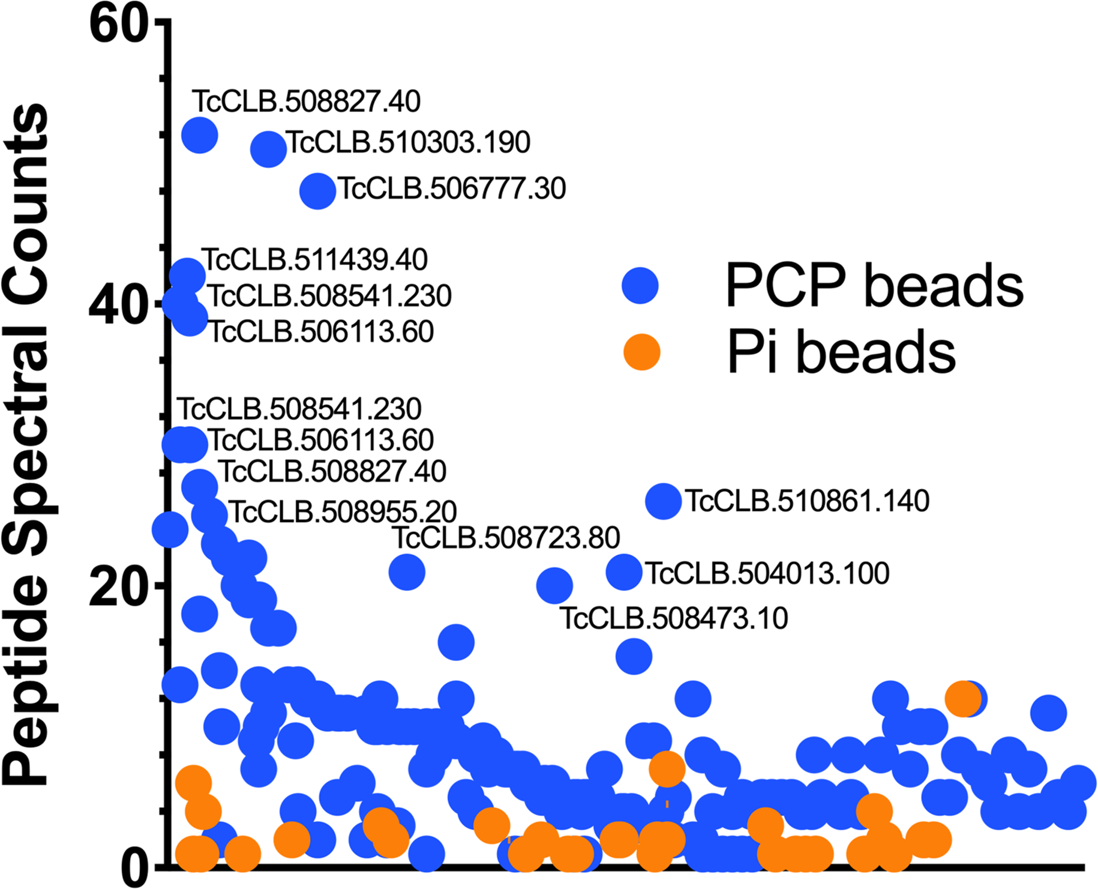
Spectral counts distribution of proteins isolated with PCP- and Pi-reagents. Scatter plot represents the peptide-spectrum matches (PSM) obtained for individual proteins following mass-spectrometric analysis of affinity precipitations from epimastigote samples under the metal–free conditions. Dots represent the means in PSM counts from three biological replicates. The most abundant protein hits are depicted with their TritrypDB accession number.

**Figure S4.**
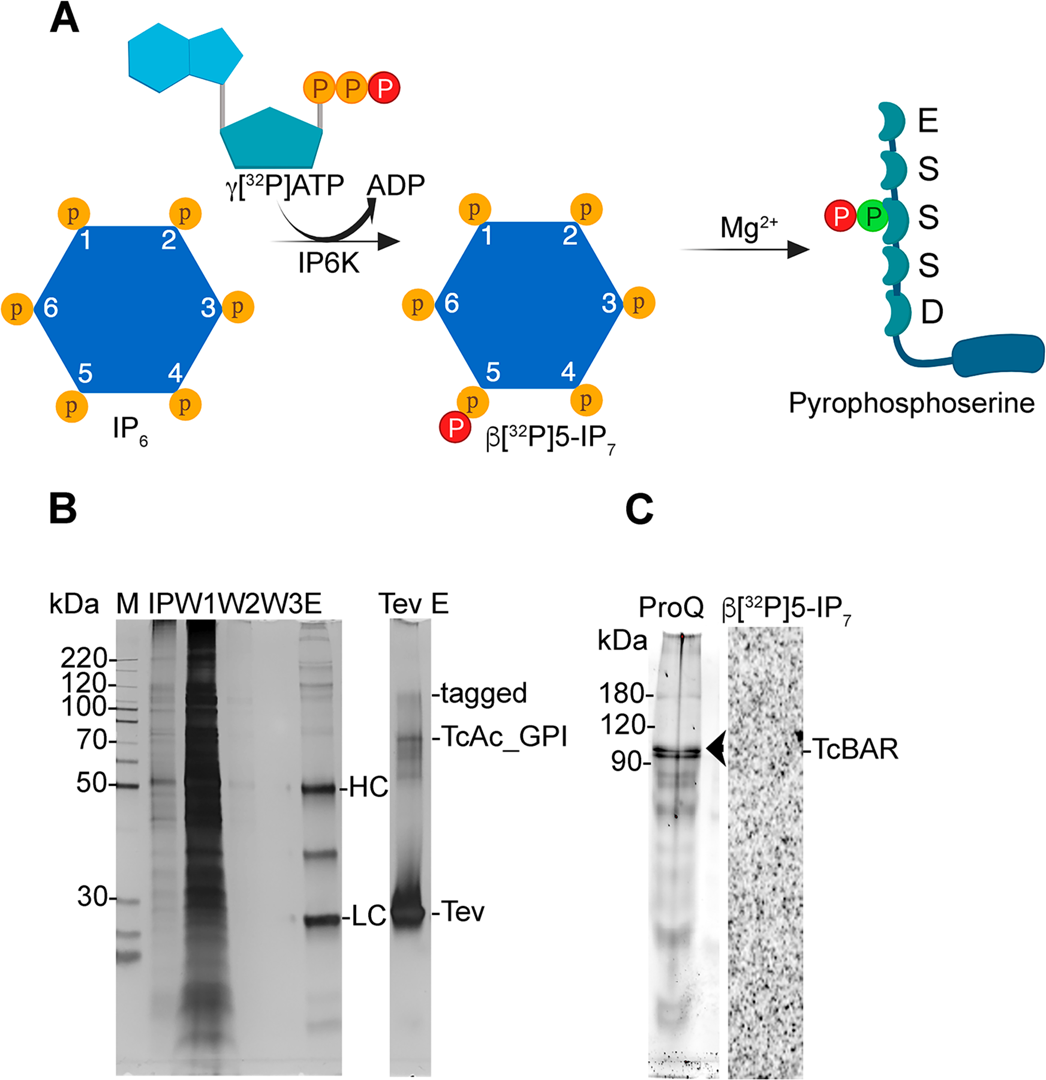
(A) Synthesis of radiolabeled 5-IP_7_ for its use in *in vitro* 5-IP_7_-phosphorylation assays. The IP_6_ kinase transfers the gamma-phosphate of [^32^P]-ATP to IP_6_ yielding [β^32^P]5-IP_7_, which in turns can transfer its β-phosphoryl group to a phosphorylated serine residue surrounded by acidic amino acids forming a pyrophosphoserine group in the target protein. (B) Immunoaffinity purification of TcAc_CPI-sm-cMyc from C-terminally tagged parasites. Parasite lysates (input, (IP) diluted 100x for gel stain) were incubated with anti-cMyc magnetic beads and after successive washings (W1-3x) protein TcAc_CPI (76 kDa) was eluted by TEV treatment (tag removed) or alkaline elution TAc_CPI-sm-cMyc (tagged = 114 kDa). The heavy (HC) and light (LC) chains of anti-cMyc were co-eluted. TEV protein size = 27 kDa. (C) Lack of pyrophosphorylation of TcBAR. C-tagged TcBAR was immunopurified from parasite lysates using anti-cMyc-magnetic dynabeads, and phosphorylation and pyrophosphorylation events were analyzed. Eluate was stained with ProQ dye (left). After incubation with [β^32^P]5-IP_7_, no signal was detected for TcBAR (*arrowhead*).

**Figure S5.**
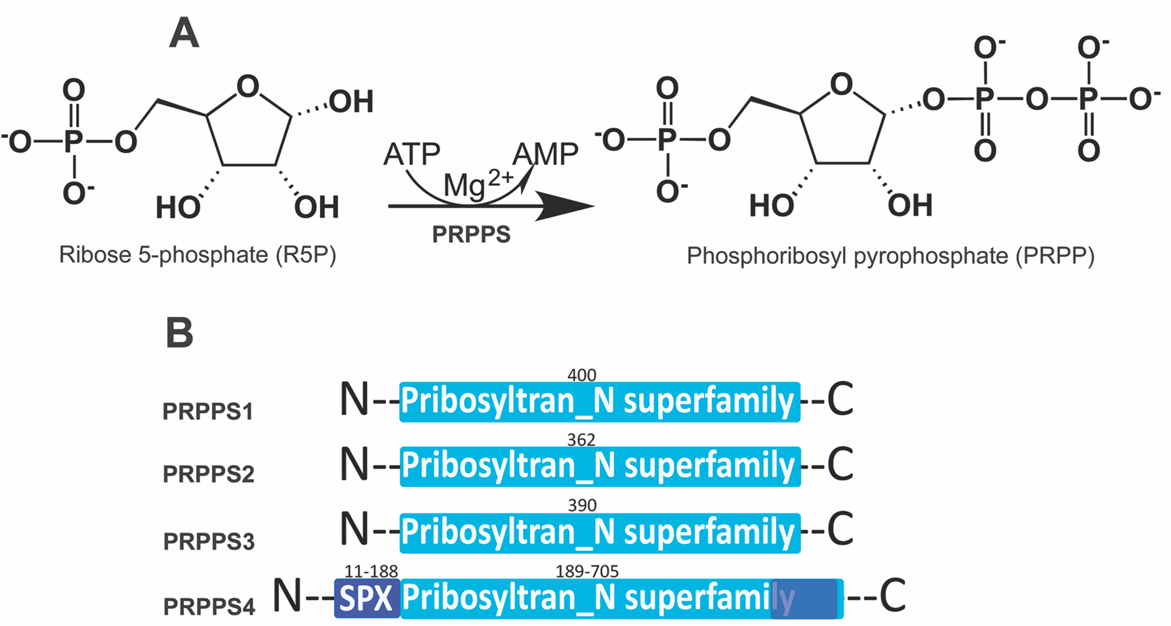
Sequence characterization of the TcPRPS4. (A) Enzymatic reaction of PRPS [EC 2.7.6.1]: a pyrophosphate group is transferred from ATP to R5P producing PRPP and AMP. (B) Protein domains identified in PRPSs of *T. cruzi*. PRPS4 is the only protein with an SPX-domain (amino acids 11-188) at the N-terminus. All of them have a phosphoribosyltransferase_N superfamily domain (Pribosyltran_N superfamily), while PRPS4 also has a region of 85 amino acids (blue square at the C-terminus), which is not found in other PRPSs.

**Figure S6.**
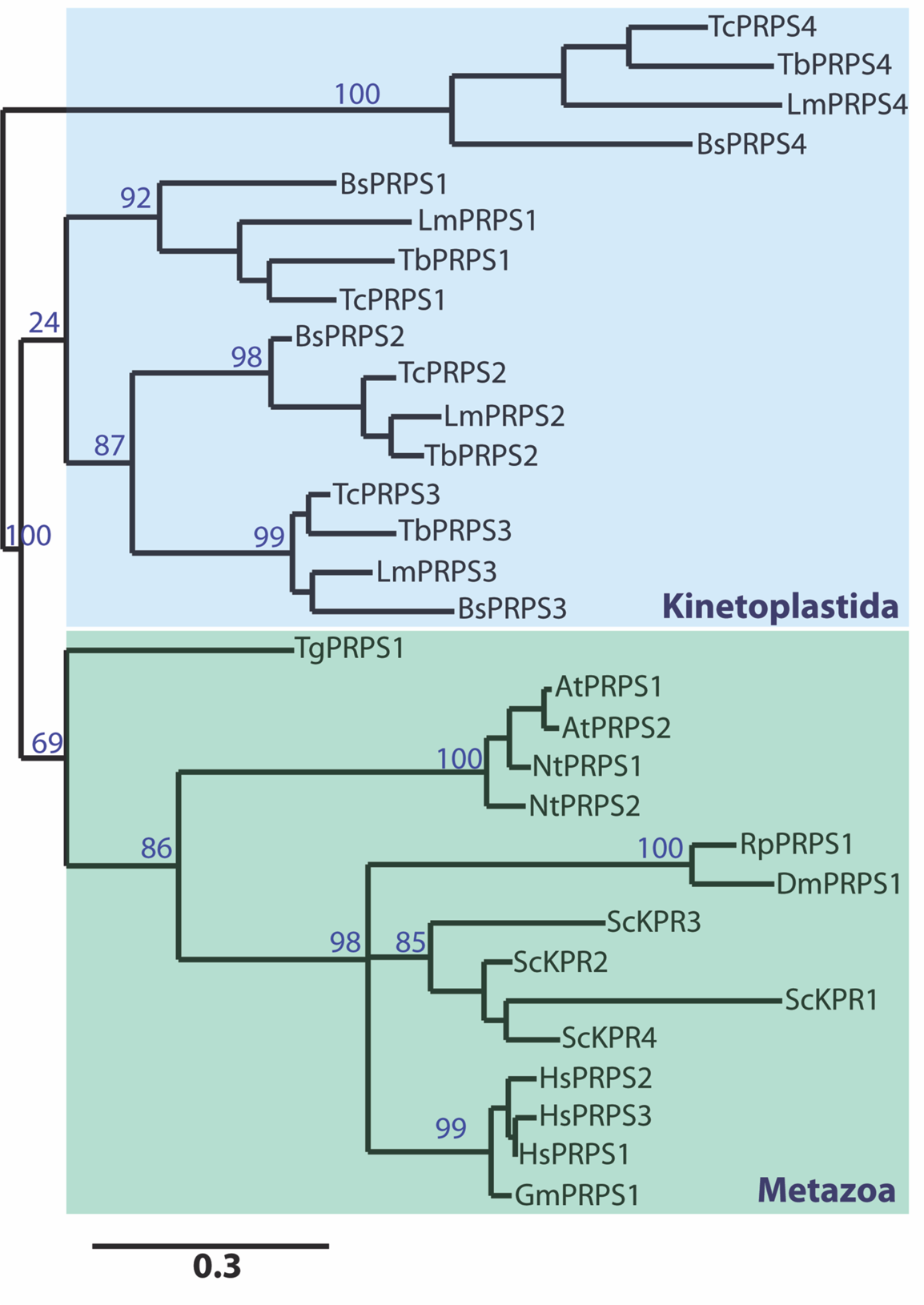
Phylogenetic tree reconstructed using the PhyloML online server (www.phylogeny.fr) with the three most common families of PRPSs found in eukaryotes. The PRPS4 group is unique and conserved in kinetoplastids showing high divergence with the other PRPSs sequences from kinetoplastids and other eukaryotes. Protein names and UniprotKb or EupathDB accession numbers are as follows: *T. cruzi* ribose-phosphate pyrophosphokinase (Q4DSS9), *T. brucei* ribose-phosphate pyrophosphokinase (Q57Z76), *Leishmania major* putative ribose-phosphate pyrophosphokinase (Q4QIB8), *Bodo saltans* ribose-phosphate pyrophosphokinase (A0A0S4IR80), *B. saltans* phosphoribosylpyrophosphate synthetase (BSAL_57745), *L. major* phosphoribosylpyrophosphate synthetase (LMJLV39_080015300), *T. brucei* phosphoribosylpyrophosphate synthetase, putative (Tb427.05.2960), (TcCLB.506321.310), *T. cruzi* phosphoribosylpyrophosphate synthetase (TcCLB.510431.250), *L. major* phosphoribosylpyrophosphate synthetase, putative (LmjF.33.1930), *T. brucei* phosphoribosylpyrophosphate synthetase, putative (Tb427.tmp.02.0530), *B. saltans* phosphoribosyl transferase, putative (BSal_54965), *T. cruzi* phosphoribosylpyrophosphate synthetase, putative (TcCLB.508717.30), *L. major* phosphoribosylpyrophosphate synthetase (LMJLV39_360066100), *T. brucei* phosphoribosylpyrophosphate synthetase, putative (Tb427.10.9430), *B. saltans* phosphoribosylpyrophosphate synthetase, putative (BSAL_71415), *Toxoplasma gondii* phosphoribosylpyrophosphate synthetase (TGARI_220100), *Arabidopsis thaliana* ribose-phosphate pyrophosphokinase 1 (Q42581), *A. thaliana* ribose-phosphate pyrophosphokinase 2 (Q42583), *Nicotiana tabacum* ribose-phosphate pyrophosphokinase 1 (A0A1S3ZNC4), *N. tabacum* ribose-phosphate pyrophosphokinase 2 (A0A1S3XTM5), *Rhodnius prolixus* putative ribose-phosphate pyrophosphokinase 1,2 (R4FNN1), *Drosophila melanogaster* phosphoribosyl pyrophosphate synthase-associated protein 2 (A0A0B4JD23), *Saccharomyces cerevisiae* ribose-phosphate pyrophosphokinase 1 (P32895), *S. cerevisiae* ribose-phosphate pyrophosphokinase 2 (P38620), *S. cerevisiae* ribose-phosphate pyrophosphokinase 3 (P38689), *S. cerevisiae* ribose-phosphate pyrophosphokinase 4 (P38063), human ribose-phosphate pyrophosphokinase 1 (P60891), human ribose-phosphate pyrophosphokinase 2 (P11908), human ribose-phosphate pyrophosphokinase 3 (P21108) and *Glossina morsitans* ribose-phosphate pyrophosphokinase (A0A1B0FPW5).

**Table S1.** Protein output obtained from epimastigote lysates enriched under metal-free conditions (EDTA^+^). Proteomic data were listed according to MASCOT score and PSM values were used for comparisons between PCP/Pi beads. Results from GO terms enrichment analysis were also included as separate worksheets.

**Table S2.** Protein output obtained from epimastigote lysates enriched in the presence of magnesium (Mg^2+^). Proteomic data were listed according to MASCOT score and PSM values were used for comparisons between PCP/Pi beads. Results from GO terms enrichment analysis were also included as separate worksheets.

**Table S3.** Protein output obtained from amastigote lysates enriched under metal-free conditions (EDTA^+^). Proteomic data were listed according to MASCOT score and PSM values were used for comparisons between PCP/Pi beads. Results from GO terms enrichment analysis were also included as separate worksheets.

**Table S4.** Protein output obtained from amastigote lysates enriched in the presence of magnesium (Mg^2+^). Proteomic data were listed according to MASCOT score and PSM values were used for comparisons between PCP/Pi beads. Results from GO terms enrichment analysis were also included as separate worksheets.

**Table S5.** Primers used in this study

