## Supplemental figures for "Protein Targets of Inositol Pyrophosphate (5-IP_7_) in the parasite *Trypanosoma cruzi*"

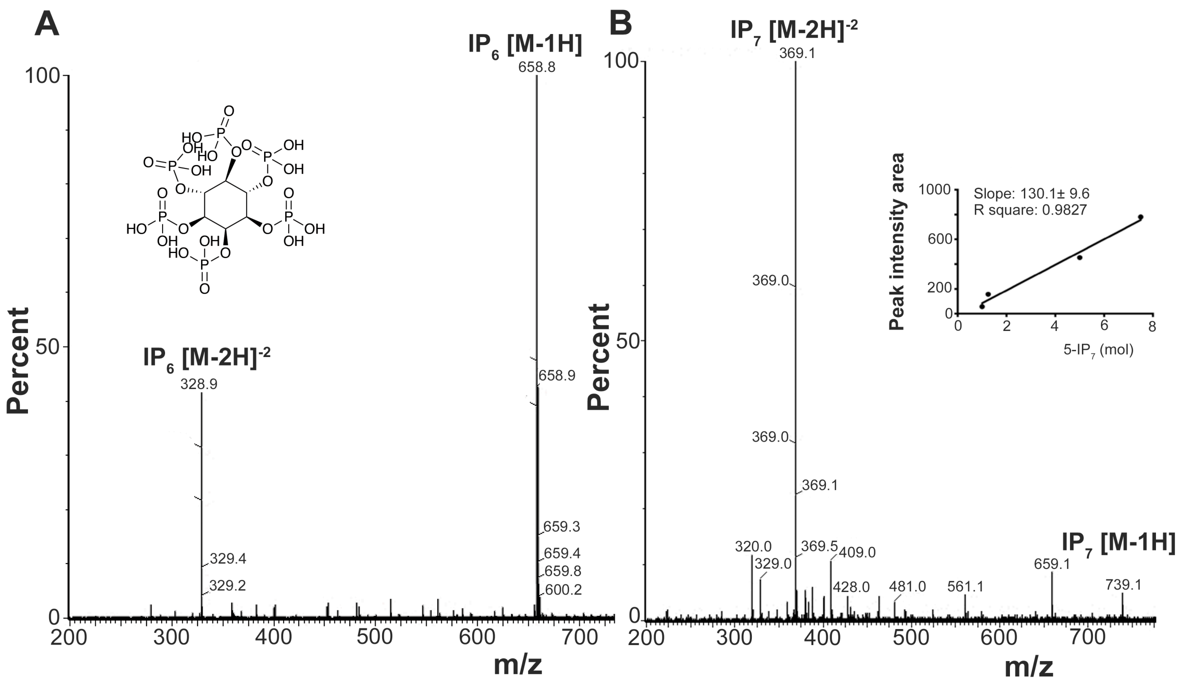


**Figure S1.** Negative ion electrospray mass spectrometry of inositol hexakisphosphate ($IP_6$) (A) and 5-diphosphoinositol-pentakisphosphate (5-$IP_7$) (B) standards. (A) Ten nmol of $IP_6$ [M = 660 Da] were injected and separated by LC. Two main ions of 329 m/z and 659 m/z (single deprotonated) are shown. Inset shows $IP_6$ structure. (B) Ten nmol 5-$IP_7$ [M = 740 Da] were injected and separated by LC. Two main ions of 369 m/z (double deprotonated) and 739 m/z (single deprotonated) are shown. Inset shows a calibration curve using 1-10 nmol of 5-$IP_7$. Peak integration area was used to determine total amount of 5-$IP_7$ in parasite extracts.

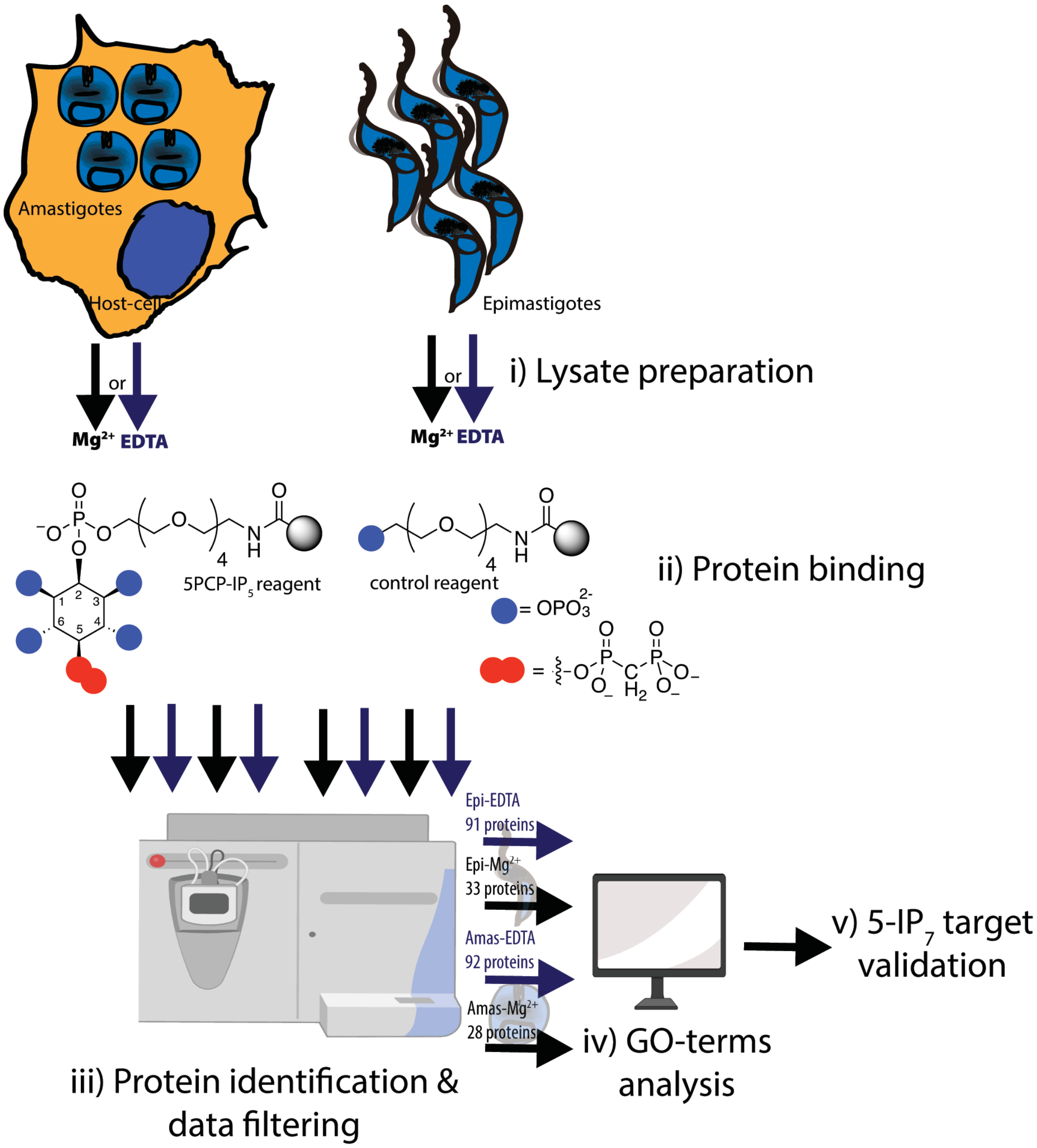


**Figure S2.** Experimental workflow followed in this study. Cell-free protein samples were prepared from proliferative stages of *T. cruzi*. Epimastigotes were grown axenically and amastigotes were obtained from gamma-irradiated Vero cells infected with cell-derived trypomastigotes. i) Lysates from both stages were prepared in the presence of magnesium ions or under the metal-free conditions. ii) Protein lysates were incubated with 5PCP-$IP_5$ or control reagent, separately. iii) Eluted proteins (from 8 different conditions) were identified by mass spectrometry and results were filtered out according to the PSM enrichment obtained in the 5PCP-$IP_5$ beads relative to phosphate beads used as control. iv) GO-terms analysis was done for all the resultant dataset hits enriched in at least two biological replicates. v) Based on protein identification data some targets were chosen for validation experiments in the parasite or *in vitro*.

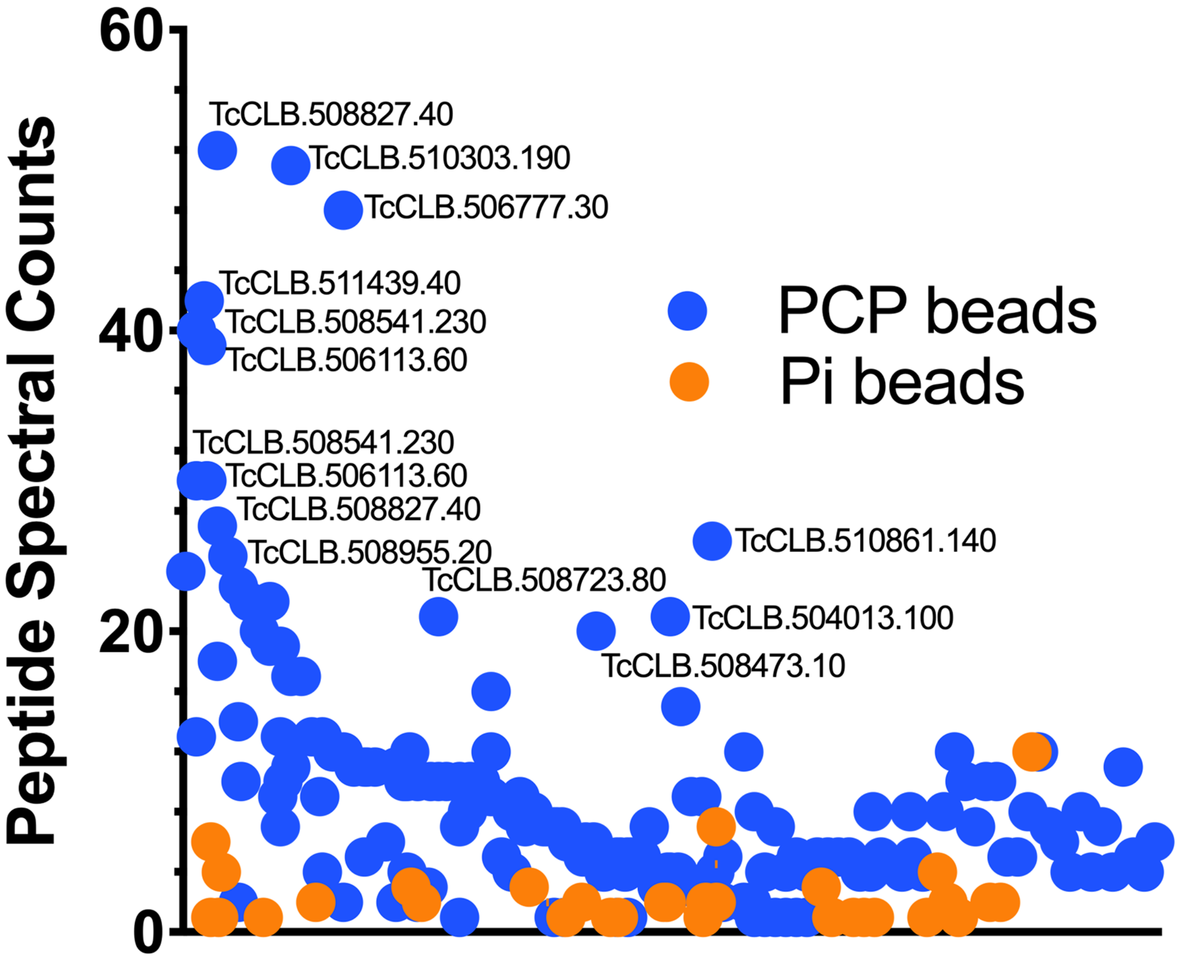


**Figure S3**. Spectral counts distribution of proteins isolated with PCP- and Pi-reagents. Scatter plot represents the peptide-spectrum matches (PSM) obtained for individual proteins following mass-spectrometric analysis of affinity precipitations from epimastigote samples under the metal–free conditions. Dots represent the means in PSM counts from three biological replicates. The most abundant protein hits are depicted with their TritrypDB accession number.

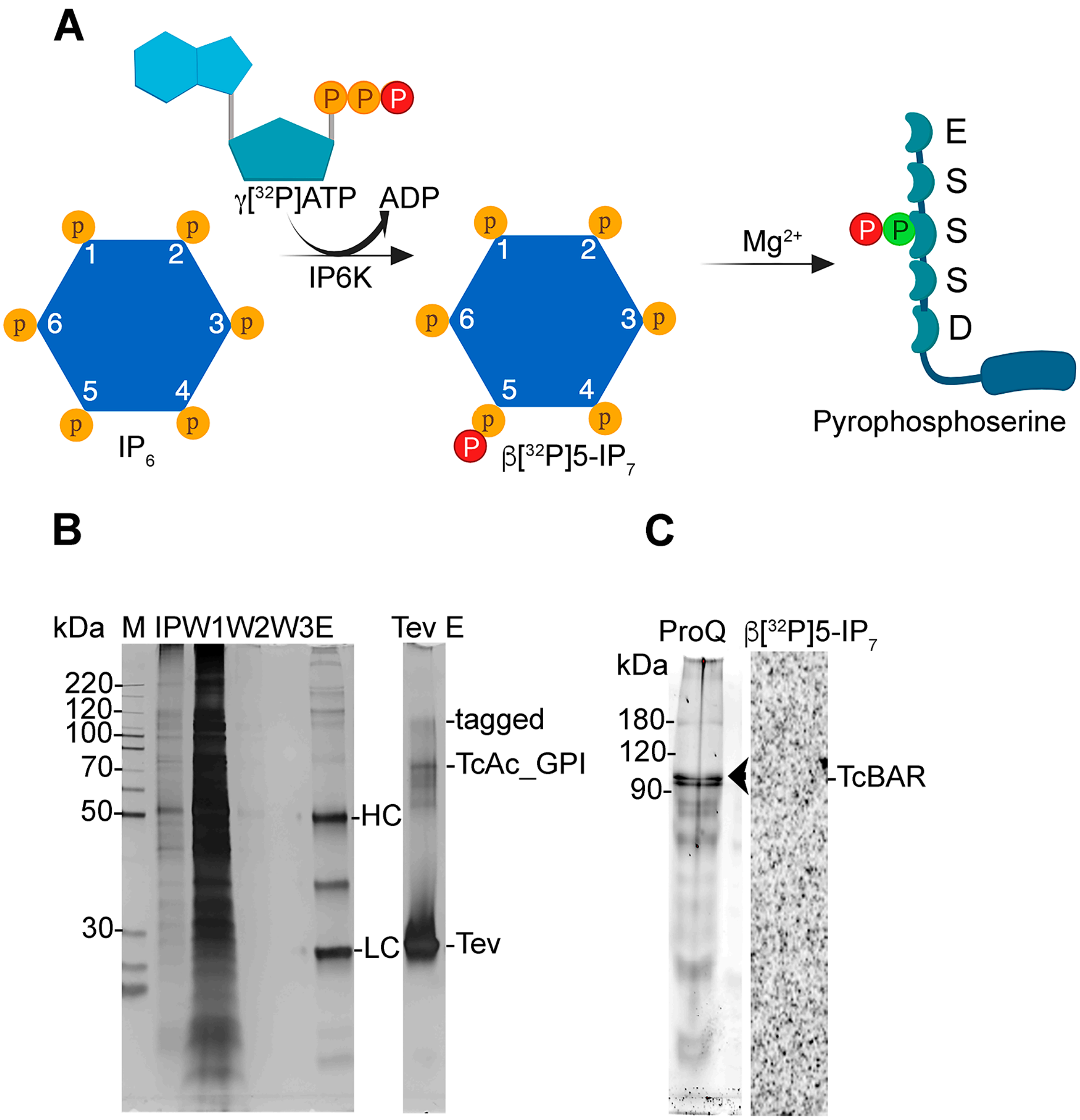


**Figure S4.** (A) Synthesis of radiolabeled 5-$IP_7$ for its use in *in vitro* 5-$IP_7$-phosphorylation assays. The $IP_6$ kinase transfers the gamma-phosphate of [$^{32}$P]-ATP to $IP_6$ yielding [$\beta^{32}$P]5-$IP_7$, which in turns can transfer its β-phosphoryl group to a phosphorylated serine residue surrounded by acidic amino acids forming a pyrophosphoserine group in the target protein. (B) Immunoaffinity purification of TcAc_CPI-sm-cMyc from C-terminally tagged parasites. Parasite lysates (input, (IP) diluted 100x for gel stain) were incubated with anti-cMyc magnetic beads and after successive washings (W1-3x) protein TcAc_CPI (76 kDa) was eluted by TEV treatment (tag removed) or alkaline elution TAc_CPI-sm-cMyc (tagged = 114 kDa). The heavy (HC) and light (LC) chains of anti-cMyc were co-eluted. TEV protein size = 27 kDa. (C) Lack of pyrophosphorylation of TcBAR. C-tagged TcBAR was immunopurified from parasite lysates using anti-cMyc-magnetic dynabeads, and phosphorylation and pyrophosphorylation events were analyzed. Eluate was stained with ProQ dye (left). After incubation with [$\beta^{32}$P]5-$IP_7$, no signal was detected for TcBAR (*arrowhead*).

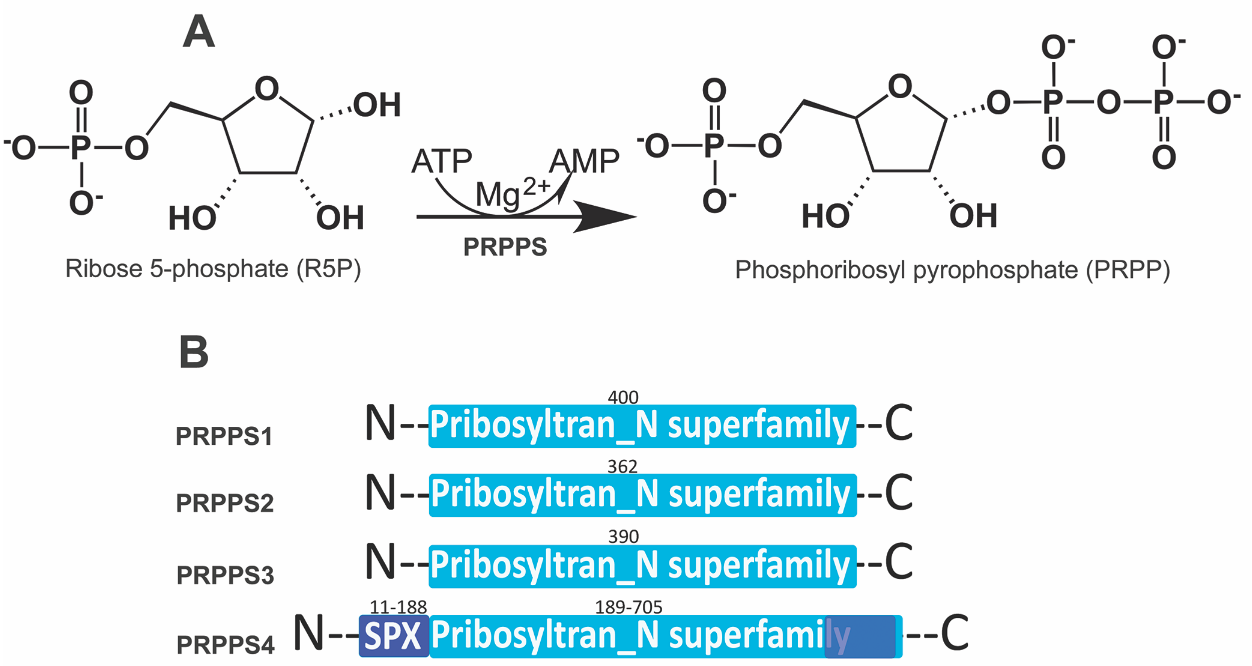


**Figure S5.** Sequence characterization of the TcPRPS4. (A) Enzymatic reaction of PRPS [EC 2.7.6.1]: a pyrophosphate group is transferred from ATP to R5P producing PRPP and AMP. (B) Protein domains identified in PRPSs of *T. cruzi*. PRPS4 is the only protein with an SPX-domain (amino acids 11-188) at the N-terminus. All of them have a phosphoribosyltransferase_N superfamily domain (Pribosyltran_N superfamily), while PRPS4 also has a region of 85 amino acids (blue square at the C-terminus), which is not found in other PRPSs.

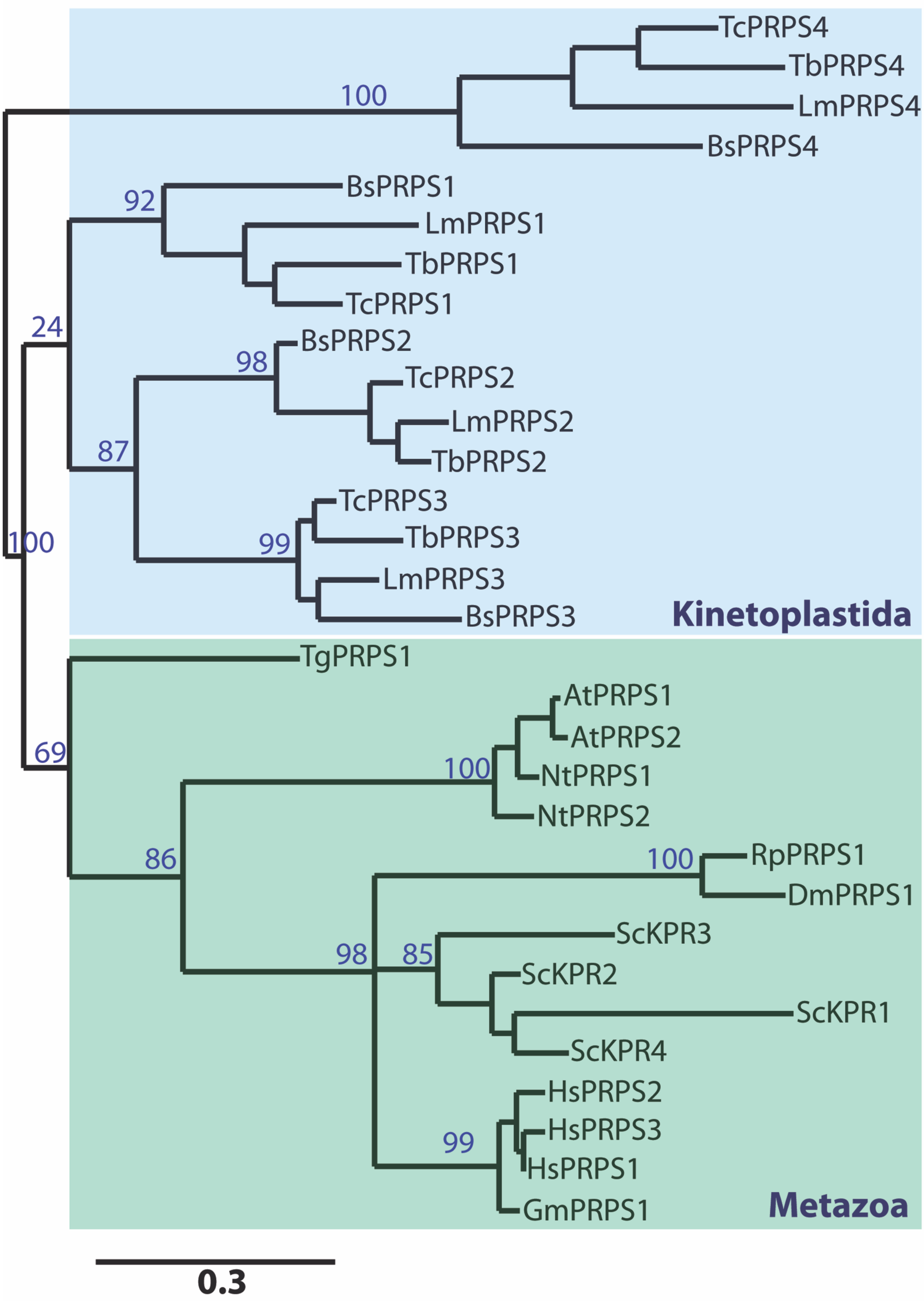
TcPRPS4
TbPRPS4
LmPRPS4
BsPRPS4
100
BsPRPS1
92
LmPRPS1
TbPRPS1
TcPRPS1
24
BsPRPS2
98
TcPRPS2
LmPRPS2
87
TbPRPS2
TcPRPS3
100
99
TbPRPS3
LmPRPS3
BsPRPS3
Kinetoplastida
TgPRPS1
AtPRPS1
AtPRPS2
69
100
NtPRPS1
NtPRPS2
86
100
RpPRPS1
DmPRPS1
ScKPR3
98
85
ScKPR2
ScKPR1
ScKPR4
HsPRPS2
HsPRPS3
99
HsPRPS1
GmPRPS1
Metazoa
0.3

**Figure S6.** Phylogenetic tree reconstructed using the PhyloML online server (www.phylogeny.fr) with the three most common families of PRPSs found in eukaryotes. The PRPS4 group is unique and conserved in kinetoplastids showing high divergence with the other PRPSs sequences from kinetoplastids and other eukaryotes. Protein names and UniprotKb or EupathDB accession numbers are as follows: *T. cruzi* ribose-phosphate pyrophosphokinase (Q4DSS9), *T. brucei* ribose-phosphate pyrophosphokinase (Q57Z76), *Leishmania major* putative ribose-phosphate pyrophosphokinase (Q4QIB8), *Bodo saltans* ribose-phosphate pyrophosphokinase (A0A0S4IR80), *B. saltans* phosphoribosylpyrophosphate synthetase (BSAL_57745), *L. major* phosphoribosylpyrophosphate synthetase (LMJLV39_080015300), *T. brucei* phosphoribosylpyrophosphate synthetase, putative (Tb427.05.2960), (TcCLB.506321.310), *T. cruzi* phosphoribosylpyrophosphate synthetase (TcCLB.510431.250), *L. major* phosphoribosylpyrophosphate synthetase, putative (LmjF.33.1930), *T. brucei* phosphoribosylpyrophosphate synthetase, putative (Tb427.tmp.02.0530), *B. saltans* phosphoribosyl transferase, putative (BSal_54965), *T. cruzi* phosphoribosylpyrophosphate synthetase, putative (TcCLB.508717.30), *L. major* phosphoribosylpyrophosphate synthetase (LMJLV39_360066100), *T. brucei* phosphoribosylpyrophosphate synthetase, putative (Tb427.10.9430), *B. saltans* phosphoribosylpyrophosphate synthetase, putative (BSAL_71415), *Toxoplasma gondii* phosphoribosylpyrophosphate synthetase (TGARI_220100), *Arabidopsis thaliana* ribose-phosphate pyrophosphokinase 1 (Q42581), *A. thaliana* ribose-phosphate pyrophosphokinase 2 (Q42583), *Nicotiana tabacum* ribose-phosphate pyrophosphokinase 1 (A0A1S3ZNC4), *N. tabacum* ribose-phosphate pyrophosphokinase 2 (A0A1S3XTM5), *Rhodnius prolixus* putative ribose-phosphate pyrophosphokinase 1,2 (R4FNN1), *Drosophila melanogaster* phosphoribosyl pyrophosphate synthase-associated protein 2 (A0A0B4JD23), *Saccharomyces cerevisiae* ribose-phosphate pyrophosphokinase 1 (P32895), *S. cerevisiae* ribose-phosphate pyrophosphokinase 2 (P38620), *S. cerevisiae* ribose-phosphate pyrophosphokinase 3 (P38689), *S. cerevisiae* ribose-phosphate pyrophosphokinase 4 (P38063), human ribose-phosphate pyrophosphokinase 1 (P60891), human ribose-phosphate pyrophosphokinase 2 (P11908), human ribose-phosphate pyrophosphokinase 3 (P21108) and *Glossina morsitans* ribose-phosphate pyrophosphokinase (A0A1B0FPW5).

**Table S1.** Protein output obtained from epimastigote lysates enriched under metal-free conditions ($EDTA^{+}$). Proteomic data were listed according to MASCOT score and PSM values were used for comparisons between PCP/Pi beads. Results from GO terms enrichment analysis were also included as separate worksheets.

**Table S2.** Protein output obtained from epimastigote lysates enriched in the presence of magnesium ($Mg^{2+}$). Proteomic data were listed according to MASCOT score and PSM values were used for comparisons between PCP/Pi beads. Results from GO terms enrichment analysis were also included as separate worksheets.

**Table S3.** Protein output obtained from amastigote lysates enriched under metal-free conditions ($EDTA^{+}$). Proteomic data were listed according to MASCOT score and PSM values were used for comparisons between PCP/Pi beads. Results from GO terms enrichment analysis were also included as separate worksheets.

**Table S4.** Protein output obtained from amastigote lysates enriched in the presence of magnesium ($Mg^{2+}$). Proteomic data were listed according to MASCOT score and PSM values were used for comparisons between PCP/Pi beads. Results from GO terms enrichment analysis were also included as separate worksheets.

**Table S5.** Primers used in this study
