## Supplementary Table S5 for "Protein Targets of Inositol Pyrophosphate (5-IP_7_) in the parasite *Trypanosoma cruzi*"

'able S5. List of oligonucleotides used in this study:

'RISPR/Cas9-mediated endogenous C-terminal tagging:

| Gene accession number | No | Oligo name | Oligonucleotide Sequence 5’-3’ | Purpose |
|---|---|---|---|---|
| TcCLB.504213.90 | 1 | sgRNA-TcIP6K | GATCGGATCCTAACGACATGACGCACCGCGGTTTTAGAGCTAGAAATAGC | Cloning of sgRNA into pTREXn for C-tagging |
| PMID: 27793988 | 2 | Rv-sgRNA-scaffold | CAGTGGATCCAAAAAAGCACCGACTCGGTG | Consensus for sgRNA amplification |
| PMID: 26199333 | 3 | Rv-HX1 | TAATTTCGCTTTCGTGCGTG | Verification of sgRNA cloning |
| TcCLB.504213.90 | 4 | Fw HR1-TcIP6K | CGAACGGGAGGAGTTGCAGCGCGGTGGGGACCCCGAGGCGACGTACGAACGGGATTTTGGGT ACATCAAAGCCCTCGATACGATGCTTACGATACTCGCCGGTACCGGGCCCCCCCTCGAG | Donor template amplification |
| TcCLB.504213.90 | 5 | Rv HR2-TcIP6K | AAAAAAGGTAATTTTGTTCTCTTCTCATGTCTTCTTCTGGGTGACGCATGTTTCCCCTCAATTGA GGCGTCTCTCTCTCTCTCTCTCTCTCCATACCTCCTGGCGGCCGCTCTAGAACTAGTGGAT | Donor template amplification |
| TcCLB.504213.90 | 6 | Fw TcIP6K_1802 | GTGTGCGGCTAGCGGG | PCR validation |
| TcCLB.504213.90 | 7 | Rv TcIP6K+193 | ACTGCAATTCGCTTGCAGTGTTTC | PCR validation |
| TcCLB.510857.10 | 8 | sgRNA-TcAc_CPI | GATCGGATCCAGAATAAACTTGAGAAGCGAGTTTTAGAGCTAGAAATAGC | Cloning of sgRNA into pTREXn for C-tagging |
| TcCLB.510857.10 | 9 | Fw HR1-TcAc_CPI | AAGTGGTGGAAGACACATATGCCAACTATATGCTGATGCATTGGAGCAGTCCCTTTATGACTTG TACCGTCTTTCAGCTGTGTTGCCTCCTCCGAAACTTGGTACCGGGCCCCCCCTCGAG | Donor template amplification |
| TcCLB.510857.10 | 10 | Rv HR2-TcAc_CPI | TGTATGAGTAATGCAGAATAGTCAAAATAATAACACAAAAAAAACAAAAGCACGCGAAAGAT CTCTCGCCCCTTCTAGGAAACAACTCATTTTCGCTTCTTGGCGGCCGCTCTAGAACTAGTGGAT | Donor template amplification |
| TcCLB.510857.10 | 11 | Fw TcAc_CPI_1902 | ACGTCCACTGAAGGGCACG | PCR validation |
| TcCLB.510857.10 | 12 | Rv TcAc_CPI+155 | TCCAACTGGCATTGTGGCAGAG | PCR validation |
| TcCLB.510515.30 | 13 | sgRNA-TcBAR | GATCGGATCCGGTGTTTGCATGCCTAATGCGTTTTAGAGCTAGAAATAGC | Cloning of sgRNA into pTREXn for C-tagging |
| TcCLB.510515.30 | 14 | Fw HR1-TcBAR | GCAGCTTCGCCCACCCACGGCGCGGGATGCACCGGATGAGCCGGTTCATTTCGAAGATGCGTT GAAAGCCACACCAGCCGATTTACAGGAACACCACATGGGTACCGGGCCCCCCCTCGAG | Donor template amplification |
| TcCLB.510515.30 | 15 | Rv HR2-TcBAR | ATGTAACATCAAAACAATGCGTCTCCAAGGAGATTTTGCGTGTACGCACTTCACACAGGTATAT TTATATATACGTATATGTGAGTGGTTGGGTGTTTGCTGGCGGCCGCTCTAGAACTAGTGGAT | Donor template amplification |
| TcCLB.510515.30 | 16 | Fw TcBAR_971 | GCGTGCTTCAGGAGCCAT | PCR validation |
| TcCLB.510515.30 | 17 | Rv TcBAR+192 | CCTCGCCATTCCCCAATGCATA | PCR validation |
| TcCLB.506211.114 | 18 | sgRNA-TcMotKin | GATCGGATCCTGGTAGCGACTTAAGAAAGTGTTTTAGAGCTAGAAATAGC | Cloning of sgRNA into pTREXn for C-tagging |
| TcCLB.506211.114 | 19 | Fw HR1-TcMotKin | AAGCTACAATTGCAGTGAAAGCAACAGTTCCCTGCAGTTTGCGTCCCGTGCCCGTGGCGTGGC GTTTGGACAAATCAAAAAGAACACCGTGGTAGCGACTGGTACCGGGCCCCCCCTCGAG | Donor template amplification |
| TcCLB.506211.114 | 20 | Rv HR2-MotKin | GTAGAAAAACATTTATCAGTTATTTTGGCGTGAGTAGACGCAGGGAATTCAAGGAAAAAAACA AGAGGCCACAGAGACACACCACCACCCCGCCTCCTACTGGCGGCCGCTCTAGAACTAGTGGAT | Donor template amplification |
| TcCLB.506211.114 | 21 | Fw TcMotKin_2323 | ATGTCCGGACAGGCGAAGG | PCR validation |
| TcCLB.506211.114 | 22 | Rv TcMotKin+151 | CTCACATTCCCTCTCGCTGACA | PCR validation |

Cloning and expression of recombinant proteins in *E. coli:*

| Gene accession number | **No** | Oligo name | Oligonucleotide Sequence 5'-3' | Purpose |
|---|---|---|---|---|
| TcCLB.507809.30 | 28 | Fw-TcSPRPS | GACGACGACAAGATGCGGACGGGCAAAGGACG | Cloning of full TcPRPS4 gene in pET32 EK/LIC |
| TcCLB.507809.30 | 29 | Fw-TcPRPS_565 | GACGACGACAAGCTGATGTCTATAGATGAAACTATTTTCAACAGCG | Cloning of TcPRPS4 without SPX sequence |
| TcCLB.507809.30 | 30 | Rv-TcSPRPS | GAGGAGAAGCCCGGTCTTCTCAAAAAGCGTGGCGAGCG | Cloning TcPRPS4 gene into pET32 EK/LIC |
| TcCLB.504213.90 | 31 | Fw-TcIP6K_Met$^{300}$-ArgRIII | GACGACGACAAGATGATGAATAGCCGCCCGACG | Cloning of ArgRIII domain from TcIP6K into pET32 EK/LIC vector |
| TcCLB.504213.90 | 32 | Rv-TcIP6K | GAGGAGAAGCCCGGTTCGGCGAGTATCGTAAGCATCGTATCG | Cloning of ArgRIII domain from TcIP6K into pET32 EK/LIC vector |
| TcCLB.510857.10 | 33 | Fw-TcAc_CPI | GACGACGACAAGATGGGTCCGCATGGCGA | Cloning of TcAc_CPI into pET32 Ek/LIC |
| TcCLB.510857.10 | 34 | Rv-TcAc_CPI | GAGGAGAAGCCCGGTTTAAAGTTTCGGAGGAGGCAACACAG | Cloning of TcAc_CPI into pET32 Ek/LIC |
| TcCLB.510857.10 | 35 | Fw-TcAc_CPI_S579D | TTTTGACTCTGATTCTCTACGAGAGC | Site-directed mutagenesis of TcAc_CPI |
| TcCLB.510857.10 | 36 | Rv-TcAc_CPI_S579D | AATAAGGCATCAACGAAAG | Site-directed mutagenesis of TcAc_CPI |
| TcCLB.510857.10 | 37 | Fw- TcAc_CPI_Ser$^{578\text{-}80\ del}$ | CTACGAGAGCTCTCATGTG | Site-directed mutagenesis of TcAc_CPI |
| TcCLB.510857.10 | 38 | Rv-TcAc_CPI_Ser$^{578\text{-}80\ del}$ | GTCAAAAAATAAGGCATCAAC | Site-directed mutagenesis of TcAc_CPI |
| TcCLB.510857.10 | 39 | Fw-TcAc_CPI_856 | GAAGGGACTTCAAAGGCAGC | Sequencing of TcAc_CPI mutants |
| TcCLB.510857.10 | 40 | Fw-TcAc_CPI_1519 | GATCAAGCAGTTGCAGCG | Sequencing of TcAc_CPI mutants |
